# Habitat specialization structures population divergence and demographic variation in Amazonian floodplain birds

**DOI:** 10.64898/2026.07.13.738219

**Authors:** Leilton Willians Luna, Gabriel Macedo, Sara Elizabeth Lipshutz, Camila Cherem Ribas, Alexandre Aleixo

## Abstract

Understanding how species respond to environmental change requires linking ecological traits to gene flow and demographic history across dynamic landscapes. Using population genomic data from twelve co-distributed Amazonian riverine island birds, we show that habitat specialization predicts genetic structure, divergence, and effective population size. Species specialized on dynamic sandbar scrub habitats exhibit higher gene flow, and higher genetic diversity despite smaller effective population sizes. These patterns are consistent with dispersal tracking shifting resources, which decouples genetic diversity from population size. In contrast, river-edge forest specialists show stronger spatial genetic structure and lower genetic diversity despite larger populations, consistent with long-term isolation in more stable habitats. We also detect regional variation in effective population sizes and genetic diversity across Amazonian river sub-basins, potentially reflecting historical differences in floodplain habitat availability. Overall, our results highlight how non-equilibrium dynamics shape genomic variation and emphasize habitat-driven dispersal in diversification and persistence in seasonal systems.

## INTRODUCTION

Evolutionary histories are shaped by how organisms respond to environmental change and these responses are mediated by species’ ecological traits. For instance, attributes such as breeding behavior, foraging strategy, and habitat specialization determine how populations interact with environmental heterogeneity, thereby influencing patterns of diversification and population size (De Kort et al., 2021; Luna et al., 2023, 2025; Miller et al., 2021; Suárez et al., 2022). As a result, co-occurring species often exhibit contrasting spatial genetic structures that reflect lineage-specific responses rather than shared environmental histories (Brüniche-Olsen et al., 2021; Luna et al., 2023; Papadopoulou & Knowles, 2016). Yet, despite the recognition of these patterns, we still lack a mechanistic understanding of how ecological traits mediate the translation of environmental structure and dynamics into evolutionary outcomes. A particularly valuable model to investigate these relationships are seasonal systems characterized by recurrent habitat turnover rather than persistent barriers.

Seasonally dynamic ecosystems, such as Amazonian floodplains, offer an opportunity to test how ecological traits mediate population divergence and demographic responses. Annual flood pulses create a heterogeneous mosaic of habitats that are repeatedly inundated and reshaped, forcing species to adjust spatial behavior in response to shifting resources, which directly constrains dispersal and population persistence (Golçalves et al., 2022; Rowedder et al., 2021; Silva et al., 2021). Besides being currently dynamic, the extent and location of Amazonian floodplains have also varied historically. During the Quaternary the availability of these habitats changed in response to variation in sea level and precipitation patterns, generating stronger and more recent population size variation in birds associated with more dynamic, early successional floodplain habitats (Sawakuchi et al. 2022), and patterns of endemism and regionalization along the apparently currently continuous floodplains (Laranjeiras et al. 2024). Thus, although these systems are often assumed to promote high connectivity relative to the adjacent non-flooded upland forests (Dalapicolla et al., 2021; Harvey et al., 2017; Jonhson et al., 2023), this view ignores substantial ecological variation within floodplains and its effects on population differentiation. Consistent with this complexity, co-occurring species frequently exhibit markedly different genetic structure (Choueri et al., 2017; Luna et al., 2023; Thom et al., 2020) and historical population sizes (Barbosa et al., 2020; Schultz et al., 2024), suggesting that interactions between habitat dynamics and species-specific traits, rather than the shared environment alone, drive evolutionary outcomes.

Here, we ask whether habitat association predicts differences in genetic divergence and demographic history among co-distributed Amazonian birds that occurs mostly in riverine islands. Amazonian river islands support bird assemblages shaped by habitat heterogeneity, ecological succession, and the continual formation and erosion of islands (Borgers et al., 2019; Remsen & Parker, 1983). Riverine island birds can be broadly classified into two ecological groups based on habitat association: river-edge forest specialists and sandbar scrub specialists (Remsen & Parker, 1983; Rosenberg, 1990). These habitat types differ in vegetation structure and stability, leading to contrasting dispersal strategies (Figure 1). River-edge forest species occupy relatively stable habitats with a developed canopy that allows vertical movement during seasonal flooding, whereas sandbar scrub specialists inhabit ephemeral, low-stature vegetation that is frequently submerged, requiring horizontal dispersal among islands to track suitable habitat (Rosenberg, 1990; Rowedder et al., 2021). We hypothesize that these contrasting habitat associations generate predictable differences in genetic and demographic patterns. Specifically, species associated with stable river-edge forests are expected to exhibit stronger genetic structure, lower contemporary gene flow, and more stable effective population sizes because they are less likely to disperse over long distances. In contrast, sandbar scrub specialists should show greater connectivity, weaker spatial genetic structure, and higher gene flow as a consequence of frequent dispersal among islands, but smaller effective population sizes due to recurrent local extinctions and episodic bottlenecks caused by habitat instability. Alternatively, if seasonal flooding homogenizes populations irrespective of habitat association, all species should exhibit similar patterns of genetic divergence and demographic history.

**Figure 1.**
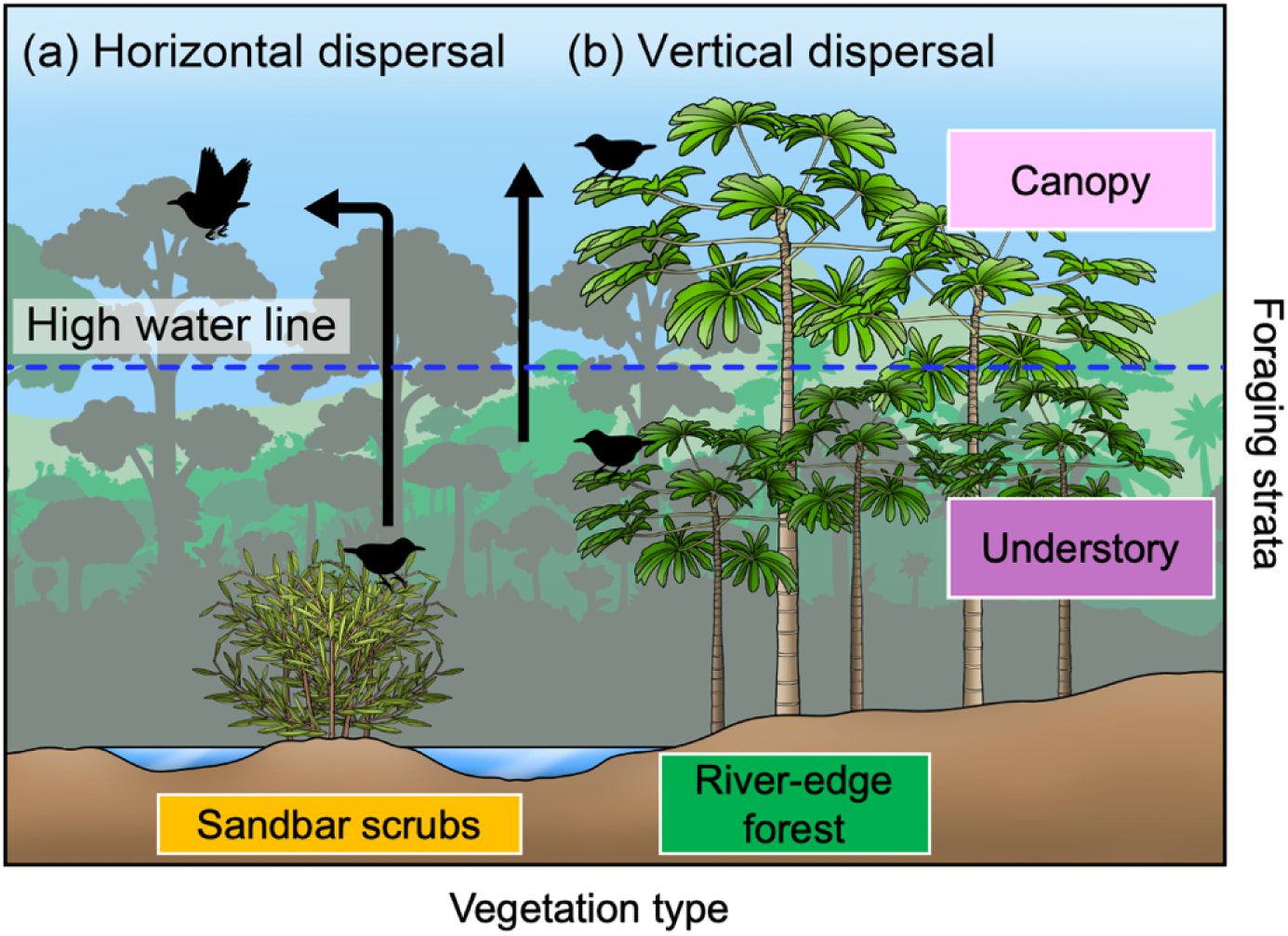
Conceptual model of habitat-driven dispersal mechanisms in Amazonian floodplains. (a) Horizontal dispersal occurs when rising water levels inundate sandbar scrub vegetation, requiring organisms to seek non-flooded areas. (b) Vertical dispersal within river-edge forests involves movement from the understory to the canopy during high-water periods. These contrasting dispersal strategies yield distinct evolutionary outcomes: horizontal dispersal is predicted to result in low genetic differentiation due to increased gene flow among adjacent localities and reduced effective population sizes due to local extinctions and episodic bottlenecks, whereas vertical dispersal should maintain high genetic differentiation due to the probability of individuals remaining in the same locality and larger effective population sizes through habitat stability. Artwork by Lucas Kías.

To test these predictions, we combined population genomic data with ecological information of species’ habitat use to conduct a comparative analysis of twelve bird species endemic to Amazonian riverine islands. We evaluated whether variation in habitat specialization explains differences in genetic structure, diversity, and demographic trajectories. We also tested whether biogeographic context contributes to spatial genetic variation, specifically whether populations within the same river sub-basin show similar patterns regardless of habitat specialization, given the regional endemism and sub-basin structuring of Amazonian floodplains (Laranjeiras et al., 2024; Luna et al., 2022; Thom et al., 2020). This framework integrates species habitat use and spatial distribution with genetic summary statistics, demographic modeling, and Bayesian phylogenetic inference, allowing us to disentangle the relative contributions of habitat use and geographic context to evolutionary responses in a dynamic environment.

## MATERIALS AND METHODS

### Study species and sampling

We sampled 12 Amazonian island bird species specialized in river-edge forests and sandbar scrubs according to Rosenberg (1990). A total of 323 samples were obtained through field expeditions, loans from ornithological collections, and public genetic databases (detailed in Supplementary Material Table S1). The number of samples per species ranged from 10 to 80 (Table 1), with an average of three samples per locality (Table S1, Figure S1). To account for shared geographic processes, we selected samples that represented the same or nearby localities across species and spanned a substantial portion of each species’ geographic range.

**Table 1.** Species included in this study, with corresponding taxonomic family, sample size used for genetic analyses, vegetation association, and primary foraging stratum. Habitat specialization classifies species as associated with either river-edge forest or sandbar scrub habitats, reflecting contrasting ecological conditions across Amazonian riverine islands.

| Family | Species | Sample size | Habitat<br>specialization<sup>1,2</sup> |
| --- | --- | --- | --- |
| Thraupidae | *Conirostrum bicolor* | 15 | River-edge forest |
| Dendrocolaptidae | *Dendroplex kienerii* | 19 | River-edge forest |
| Furnariidae | *Mazaria propinqua* | 28 | Sandbar scrubs |
| Furnariidae | *Cranioleuca vulpecula* | 26 | Sandbar scrubs |
| Furnariidae | *Furnarius minor* | 10 | Sandbar scrubs |
| Tyrannidae | *Serpophaga hypoleuca* | 16 | Sandbar scrubs |
| Tyrannidae | *Stigmatura napensis* | 29 | Sandbar scrubs |
| Thamnophilidae | *Myrmochanes hemileucus* | 21 | Sandbar scrubs |
| Thamnophilidae | *Myrmotherula klagesi* | 19 | River-edge forest |
| Thamnophilidae | *Myrmotherula assimilis* | 38 | River-edge forest |
| Thamnophilidae | *Thamnophilus cryptoleucus* | 22 | River-edge forest |
| Thamnophilidae | *Myrmoborus lugubris* | 80 | River-edge forest |

Ecological classifications were compiled from published sources: <sup>1</sup>Rosenberg 1990; <sup>2</sup>Remsen & Parker 1983.

### Species habitat preferences

We compiled information on bird habitat specialization from published sources. Habitat specialization was classified following the systems proposed by Remsen and Parker (1983) and Rosenberg (1990), which are based on standardized field observations quantifying the frequency of occurrence of each species across vegetation types. Species were assigned to one of two habitat categories regarding their habitat use within riverine islands: river-edge forest specialists or sandbar scrub specialists (Table 1). River-edge forests are relatively stable habitats dominated by short and tall stands of *Cecropia* trees (typically 20–25 m in height) that are not completely submerged in the high-water season by seasonal flooding and provide a well-developed canopy. In contrast, sandbar scrub habitats are ephemeral environments dominated by low (1–2 m) shrub vegetation, primarily of the genera *Tessaria* and *Salix*, that are frequently reshaped or submerged during seasonal flood cycles (Junk et al., 2011, 2012).

### Genomic data processing

For all bird species, we obtained genomic data using a probe set for 2,321 ultraconserved elements (UCEs) and 96 exons. UCE’s flanking regions and exon’s third codon positions have been shown to contain variability with reliable statistical resolution for inferences about recent phylogeographic patterns and population history (Amaral et al., 2018; Luna et al., 2022, 2023; Thom et al., 2020). DNA from all samples was extracted with the QIAGEN DNeasy tissue and blood kit (Valencia, CA, USA) following the manufacturer’s protocol. Subsequent laboratory procedures were performed by RAPiD Genomics (Gainesville, FL, USA). Raw paired-end reads of 150bp were obtained through automatic sequencing in the Illumina Hiseq 2500. Sequence quality control and reads count were evaluated in FastQC 0.11.5 (Andrews 2014). Sequences were then demultiplexed and the reads from low-quality adapters and bases were removed (Phred quality < 30, read length > 150 bp) using the Illumiprocessor 2.0 (Faircloth et al. 2012). The clean reads in contigs were assembled in Trinity 2.4 (Grabherr et al. 2011), using the default value of the nucleotide sequence size (kmer = 25). A set of 2,321 UCEs and 96 exons were mapped and identified using phyluce_assembly_match_contigs_to_probes implemented in PHYLUCE 1.4 (Faircloth et al. 2016). Incomplete loci matrices were generated for each species (including missing loci for some samples), establishing a completeness value of 0.8. We used BLAST 2.7.7 (Camacho et al. 2009) against zebra finch reference genome (bTaeGut7.mat; NCBI RefSeq. GCF_048771995.1) to identify and remove loci linked to the sexual chromosome Z, to avoid the inclusion of markers with different ploidy. Finally, all sequences were aligned in MAFFT 7.4 (Koyoh & Standley 2013).

For SNPs calling, we constructed a matrix including all individuals without filtering missing-data and trimming, where the longest sequences were selected as references. The reference sequences were aligned with the matrix of reads for each sample using the BWA 0.7.1 (Li & Durbin, 2009). The aligned reads with the references were mapped using a hard-masking low-quality criterion (< 30), maintaining sites with the minimum read depth of > 10, using the Unified Genotyper and Variant Annotator of GATK 3.8.1 (McKenna et al., 2010). The final matrix was assembled by randomly calling one SNP per locus from a database of biallelic SNPs without missing data, using a minor allele frequency (MAF) of 0.01.

### Population genetic structure

We determined the number of populations (*K*) that best represent the current pattern of genetic structure within each taxon using conStruct 1.0.4 in R (Bradburd et al., 2018). This method estimates the ancestry proportion for each sample at a given *K* by modeling genetic variation as a two-dimensional set of population layers and quantifying how genetic relationships among individuals decrease with distance. To assess the impact of isolation by distance on genetic structure, we applied two types of models: one accounting for spatial distance between samples (spatial model) and another ignoring the effect of distance (non-spatial model). We tested values of *K* ranging from 1 to 6, corresponding to the maximum number of Amazonian River sub-basins occupied by the studied species.

Model selection was performed to identify the value of *K* that best fit the data. We used cross-validation to evaluate predictive accuracy, training the models with 90% of the data to estimate the posterior distribution of parameters and calculating the log-likelihood for the remaining 10% test partition to determine the posterior mean value for each *K*. To exclude spurious *K* values, we calculated the relative contribution of each simulated *K* to the total covariance. *K* values contributing less than 0.02 to the total covariance were excluded following Bradburd et al. (2018). Each simulation was replicated 10 times, with 50,000 Markov chain Monte Carlo iterations per run.

To assess genetic–geographic covariation within each taxon, we performed Procrustes analysis using the R package vegan 2.5-7 (Oksanen et al., 2013). PCA analysis was conducted on SNP matrices with the ‘dudi.pca’ function from adegenet 2.1.3 R package (Jombart, 2008), summarizing genetic variation in the first two PCs. We then used the ‘protest’ function from vegan 2.5-7 to compare genetic and geographic similarity, yielding the Procrustes similarity statistic *t*, ranging from 0 (low similarity) to 1 (high similarity). Significance was assessed via 10,000 permutations of sample coordinates.

### Population differentiation and genetic diversity

We quantified genetic differentiation and compared within-population genetic diversity among pairs of genetically distinct populations within each species. Genetic differentiation was estimated using Weir and Cockerham’s weighted *F*_ST_, implemented in the R package hierfstat v0.5-11 (Goudet, 2005). This estimator corrects for sampling variance in allele frequency estimates and is robust to unequal and small sample sizes, making it appropriate for comparisons across heterogeneous spatial scales (Weir & Cockerham, 1984). Genetic diversity within populations was assessed by calculating nucleotide diversity (π), observed heterozygosity (Ho), and Tajima’s *D* using VCFtools v4.1 (Danecek et al., 2011). These complementary metrics capture different aspects of population genetic variation, with π and Ho reflecting overall levels of genetic diversity and Tajima’s *D* providing insight into departures from neutral expectations associated with demographic processes such as population expansion or bottlenecks (Tajima, 1989).

### Demographic models

To assess how habitat specialization influences species’ demographic history, we used a coalescent model-based approach implemented in Fastsimcoal 2.5.2 (Excoffier et al., 2013). Our goal was to test scenarios of population divergence either in isolation (without recent gene flow) or with secondary contact (with recent gene flow). We estimated parameters such as divergence time, migration rate, and effective population size. Since *Furnarius minor* exhibited no population structure (*K* = 1), we estimated its effective population size using a model that assumes no migration or divergence time. To do this, we generated joint site frequency spectra (jSFS) from SNP variants for each species using easySFS 0.0.1 (https://github.com/isaacovercast/easySFS; Gutenkunst et al., 2009). We ran 100,000 simulations with 100 independent replicates per model and employed an information theory approach for model selection (Burnham & Anderson, 2003), estimating AIC, relative likelihood, and AIC weights to select the best-fitting model for each species. For the selected model, we performed 50 parametric bootstraps to obtain mean estimates and 95% confidence intervals for each parameter. We used a mutation rate of 2.5×10^-9^ substitutions/site/generation, based on the genome-wide average of other well-studied passerine lineages (Nadachowska-Brzyska et al., 2015). To calculate the relative times of population divergence events, we used a generation length of 2 years, an estimated average value for Passeriformes derived from models incorporating the age of first reproduction (Bird et al., 2020). In Fastsimcoal2, per-generation migration rates (m) are estimated bidirectionally—that is, from population A to B and from B to A. To calculate the average migration rate between populations, we summed both directional estimates and divided them by two.

### Bayesian phylogenetic modeling

We analyzed the metrics of genetic diversity, differentiation, and demographic history with Bayesian phylogenetic multilevel models using brms 2.23.0 (Bürkner, 2017; Bürkner et al., 2024) and bayestestR 0.17.0 (Makowski et al., 2019). We used the inverse phylogenetic covariation matrix as a group-level effect to control for evolutionary relationships of the species (Hadfield, 2024). We estimated the phylogenetic relationships of the species of interest with RAxML 8.2.12 (Stamatakis, 2014) based on UCEs and exons sequence data. Sequence alignments were generated with MAFFT 7.4 (Katoh & Standley, 2013), and poorly aligned regions were excluded using trimAl 1.4 (Capella-Gutiérrez et al., 2009) with gap threshold of 0.2. Phylogenetic inference was conducted using the GTR+G model with 1,000 bootstrap replicates to assess node support. A maximum credible clade tree was created using TreeAnnotator 1.8.4 (Drummond & Rambaut, 2007). Additionally, because the species had repeated measures corresponding to each sampled population, we also specified population locations as a group-level effect. As response variables, we used the metrics of genetic diversity and differentiation (*F*_ST_, genetic-geographic similarity – *t_0_*, nucleotide diversity – π, observed heterozygosity – Ho, Tajima’s *D*) and demographic history (effective population size – Ne, migration rate, divergence time, time of secondary contact). As explanatory variables, we specified habitat specialization and the hydrographic sub-basin of the populations. We used the R package emmeans 2.0.1 (Lenth et al., 2025) to obtain contrasts and credible intervals of the genetic metrics between sub-basins. We centered and scaled all response variables to a mean of zero and a standard deviation of 1. We specified weakly informative normal priors for explanatory variables (N ∼ 0, 1) and a skew normal distribution for response variables to account for skewness in the response variables. We evaluated model convergence and fit using the Gelman-Rubin diagnostic (Gelman & Rubin, 1992), posterior predictive checks, and chain autocorrelation plots. We ran all models for 10,000 iterations (with half used in burn-in), in four Markov chains, and a thinning rate of one (Link & Eaton, 2012). We conducted all statistical analyses in R 4.5.0.

## RESULTS

### Sequence statistics data

Our sequence alignment recovered an average of 2,351 loci per species, about 97% of 2,417 loci in the total UCEs and exons probe set. The average number of informative loci per species was 15,106, with an average of 9,436 SNPs for each species. When we selected one biallelic SNP per locus with no missing-data and MAF of 0.01, we recovered between 708 to 1,555 SNPs, with an average of 1,186 SNPs per species (Table S2).

### Population genetic structure

Within-species analyses revealed discrete genetic structure ranging from *K* = 1 to 3 (Figures S2, S3). Figure 2 illustrates this pattern using two representative species, *F. minor* and *Myrmotherula klagesi* with contrasting habitat specialization. Spatial models generally outperformed non-spatial models in describing genetic patterns (Table S3-S14; Figure S2), with the exception of *Mazaria propinqua* (Figure S2c). Cross-validation and low marginal covariation (< 0.02) supported *K* values between 1 and 3 across all twelve species (Tables S3–S14; Figure S2). Only *Furnarius minor* was best described by *K* = 1, consistent with a continuous cline of genomic variation rather than discrete population structure.

**Figure 2.**
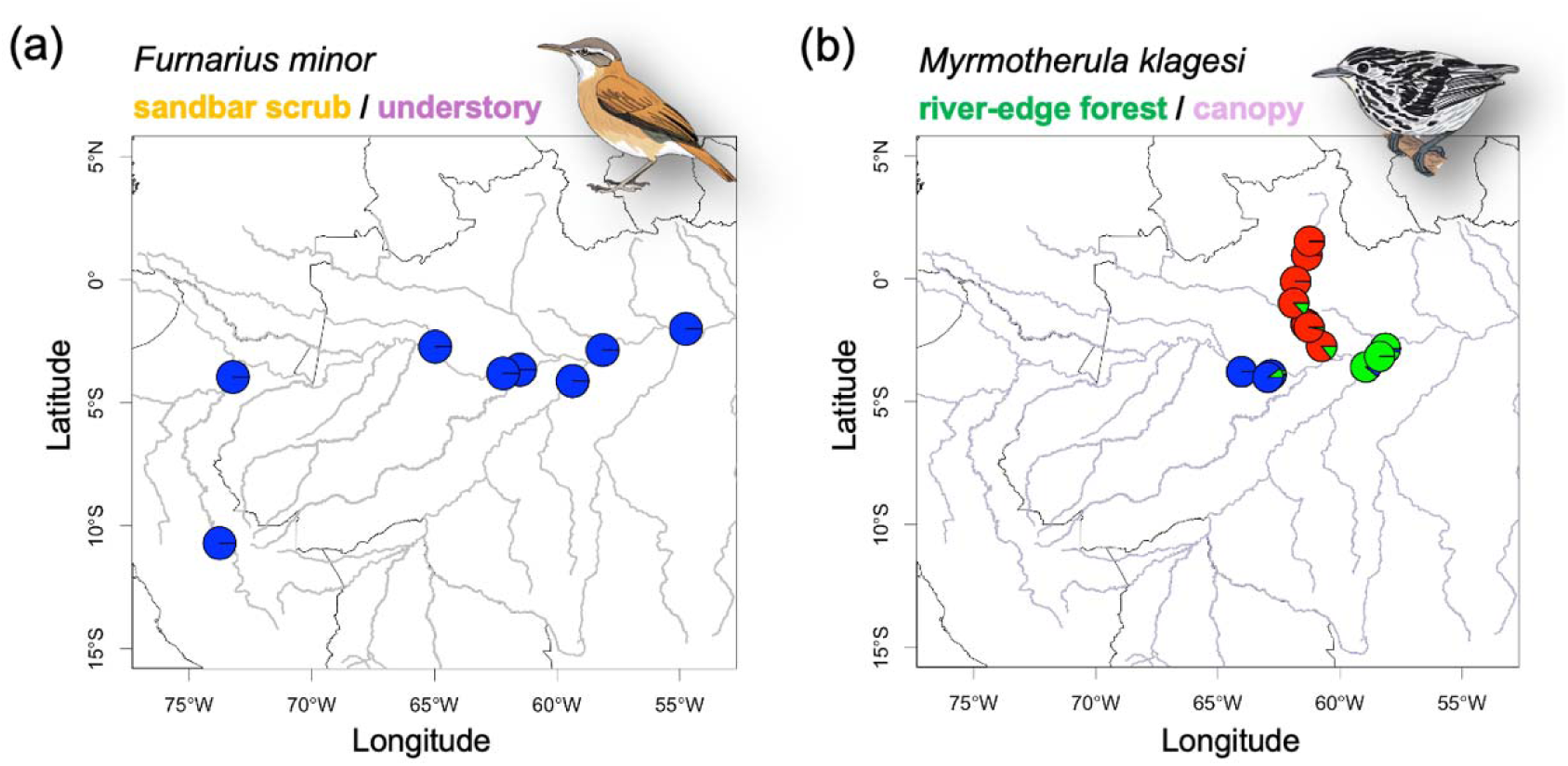
Spatial genetic structure in two Amazonian riverine bird species with contrasting habitat specialization. Population structure plots illustrate divergent patterns between species: (a) *Furnarius minor*, a specialist of the ground and understory strata of *Tessaria* vegetation on ephemeral sandbar islands, shows no population subdivision; whereas (b) *Myrmotherula klagesi*, a canopy specialist associated with *Cecropia* river-edge forests on islands and margins, exhibits three genetically differentiated populations with small admixture proportion. Bird artwork by Leilton W. Luna.

Procrustes analyses detected significant associations between genetic and geographic distances in all species (*t□*, *p* < 0.001), indicating pervasive spatial autocorrelation, although its strength varied substantially (*t□*= 0.575–0.885; Table 1; Figure S3). Spatial autocorrelation was consistently weaker in sandbar scrub specialists (mean *t□*= 0.67 ± 0.08) than in river-edge forest species (mean *t*□= 0.78 ± 0.06), indicating reduced spatial genetic structure in species occupying more dynamic habitats.

### Genetic differentiation and diversity

Genetic differentiation among populations within species ranged from low to moderate, with pairwise *F*_ST_ values spanning 0.03 to 0.217 (Table S15). Nucleotide diversity varied substantially among the population identified in the population structure analysis, from π = 0.00138 in *Myrmotherula assimilis* to π = 0.00266 in *Myrmochanes hemileucus*, and observed heterozygosity ranged from 0.0716 in *Myrmoborus lugubris* to 0.2524 in *Myrmochanes hemileucus* (Table S16). Tajima’s *D* values were heterogeneous across populations, including both negative and positive estimates, consistent with mixed demographic histories involving expansions and recent contractions (Table S16). Broadly, river-edge forest species exhibited higher genetic differentiation (mean *F*_ST_ = 0.121), lower genetic diversity (mean π = 0.0017; mean Ho = 0.141), and predominantly negative Tajima’s *D* values, whereas sandbar scrub species showed lower differentiation (mean *F*_ST_ = 0.059), higher genetic diversity (mean π = 0.0021; mean Ho = 0.182), and mostly positive Tajima’s *D* values (Tables S15, S16).

### Demographic models

Demographic model comparisons supported recent gene flow in most species, with only *Myrmotherula assimilis* showing strict isolation (Table 2). Across taxa, divergence times spanned ∼2,200 to 257,000 years, effective population sizes varied widely (∼2,400 to >300,000), and secondary contact was generally recent (890–65,000 years ago), with migration rates showing strong asymmetry among populations (0.221–6.036; Tables S17–S28). Broadly, river-edge forest species were characterized by larger effective population sizes (mean Ne = 107,750), older divergence times (mean tdiv = 107,638 years) and secondary contact (mean Tsc = 29,134 years), and lower migration rates (mean m = 1.37), whereas sandbar scrub species showed smaller Ne (mean = 22,262), more recent divergence (mean tdiv = 13,254 years) and contact (mean Tsc = 4,638 years), and higher migration rates (mean m = 2.181; Tables S17–S28).

**Table 2.**
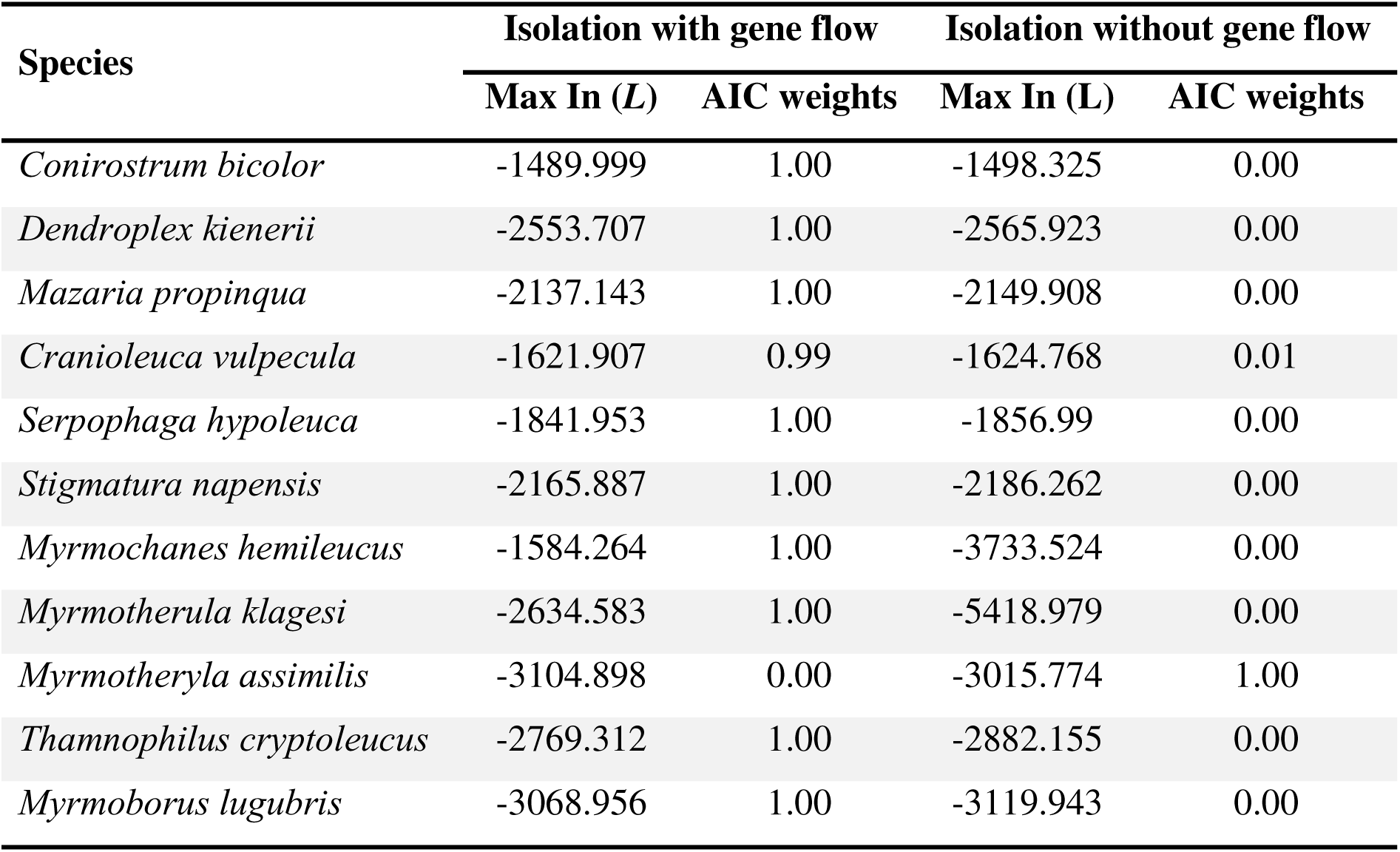
Model selection comparing demographic scenarios with and without gene flow inferred using Fastsimcoal2 for species exhibiting population subdivision. For each species, maximum composite likelihood (Max ln(L)) and Akaike Information Criterion weights (AIC weights) are reported; higher AIC weights indicate stronger model support. *Furnarius minor* was excluded due to lack of population subdivision.

| Species | Isolation with gene flow |  | Isolation without gene flow |  |
| --- | --- | --- | --- | --- |
|  | Max In (L) | AIC weights | Max In (L) | AIC weights |
| *Conirostrum bicolor* | -1489.999 | 1.00 | -1498.325 | 0.00 |
| *Dendroplex kienerii* | -2553.707 | 1.00 | -2565.923 | 0.00 |
| *Mazaria propinqua* | -2137.143 | 1.00 | -2149.908 | 0.00 |
| *Cranioleuca vulpecula* | -1621.907 | 0.99 | -1624.768 | 0.01 |
| *Serpophaga hypoleuca* | -1841.953 | 1.00 | -1856.99 | 0.00 |
| *Stigmatura napensis* | -2165.887 | 1.00 | -2186.262 | 0.00 |
| *Myrmochanes hemileucus* | -1584.264 | 1.00 | -3733.524 | 0.00 |
| *Myrmotherula klagesi* | -2634.583 | 1.00 | -5418.979 | 0.00 |
| *Myrmotheryla assimilis* | -3104.898 | 0.00 | -3015.774 | 1.00 |
| *Thamnophilus cryptoleucus* | -2769.312 | 1.00 | -2882.155 | 0.00 |
| *Myrmoborus lugubris* | -3068.956 | 1.00 | -3119.943 | 0.00 |

### Comparative population genetics analyses

Habitat type strongly structured genomic variation and demographic history across species (Figure 3). Sandbar scrub specialists consistently exhibited higher connectivity and weaker spatial genetic structure than river-edge forest species, with markedly lower genetic differentiation (*F*_ST_: β = −0.77, 95% CI: −1.28 to −0.31, pMPE < 0.001; Table S36) and higher migration rates (β = 1.02, 95% CI: 0.49 to 1.54, pMPE < 0.001; Table S37), indicating extensive gene flow among populations. These species also displayed shorter divergence times among populations (β = −0.45, 95% CI: −0.95 to −0.05, pMPE = 0.03; Table S38) and shorter times since populations secondary contact (β = −0.57, 95% CI: −1.06 to −0.16, pMPE < 0.01; Table S39), suggesting recent and dynamic population histories. Despite lower effective population sizes (Ne: β = −0.56, 95% CI: −1.06 to −0.08, pMPE = 0.02; Table S29), sandbar scrub species maintained high genetic diversity, including greater nucleotide diversity (β = 1.22, 95% CI: 0.72 to 1.71, pMPE = 0.01; Table S31), elevated observed heterozygosity (β = 1.40, 95% CI: 0.84 to 1.93, pMPE < 0.001; Table S33), higher Tajima’s *D* (β = 1.08, 95% CI: 0.61 to 1.54, pMPE < 0.001; Table S35), and weaker isolation by distance (t_0_: β = −0.87, 95% CI: −1.25 to −0.45, pMPE < 0.001; Table S40). Collectively, these patterns indicate that sandbar scrub specialists maintain high within-species diversity while experiencing frequent gene flow among populations and reduced long-term population isolation.

**Figure 3.**
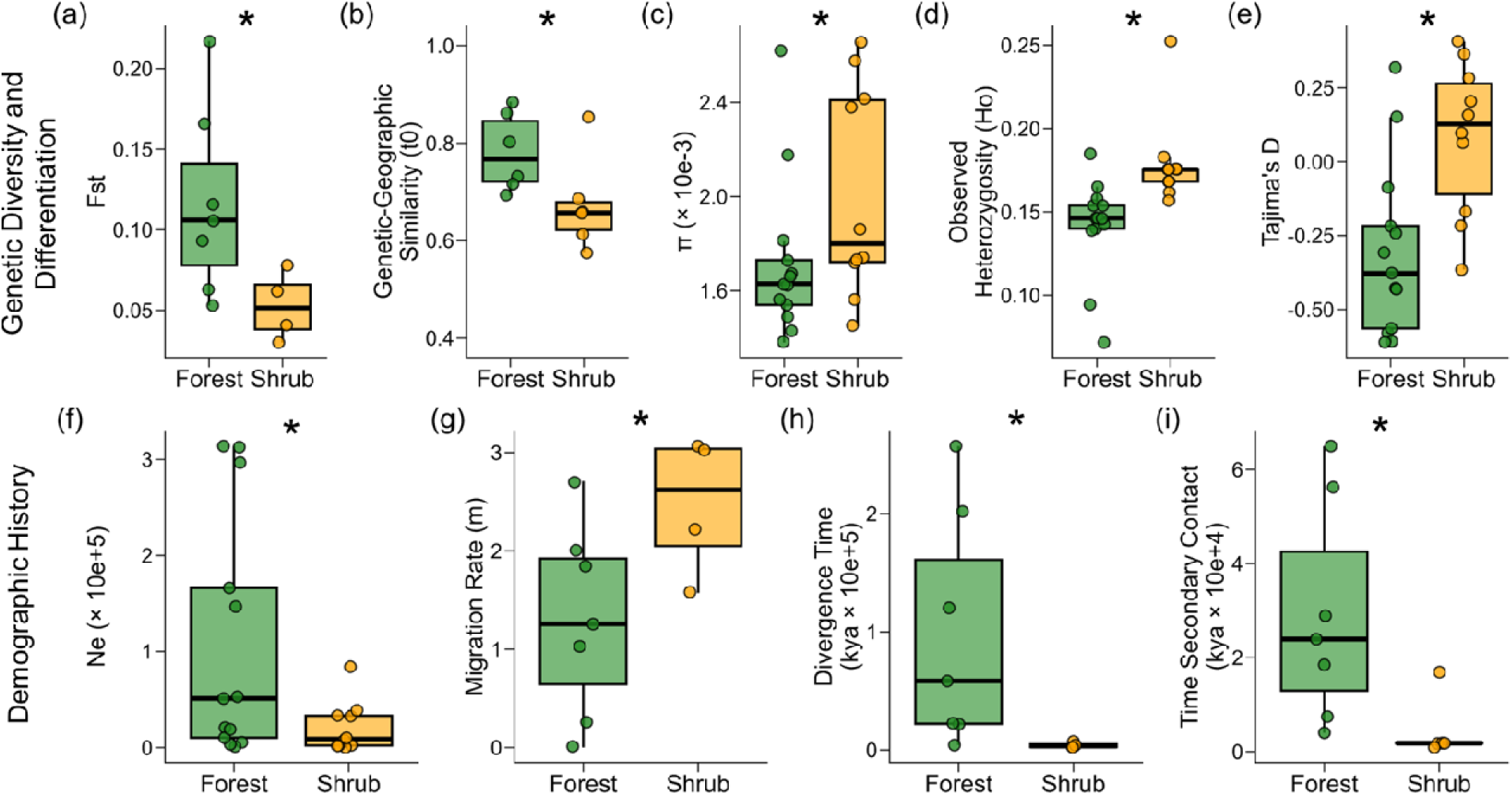
Habitat type predicts population genetic diversity, differentiation, and demographic history in an Amazonian riverine bird community. Boxplots compare populations associated with river-edge forest (green) and sandbar shrub habitats (orange). Genetic diversity and differentiation metrics include (a) genetic differentiation (*F*_ST_) between population pairs within species, (b) genetic–geographic similarity (*t*□) pattern of isolation-by-distance, (c) within population nucleotide diversity (π), (d) observed heterozygosity (H□), and (e) Tajima’s *D*. Demographic parameters inferred from the best-fit model in Fastsimcoal2 include (f) current effective population size (Ne), (g) migration rate, (h) divergence time, and (i) time of secondary contact. Boxplots follow Tukey’s convention (median, interquartile range, and 1.5× IQR whiskers), with points representing population-level estimates. Asterisks denote comparisons where 95% confidence intervals do not overlap zero.

Hydrological sub-basins accounted for relatively little variation (Figure 4). Nucleotide diversity was significantly lower in the Branco and Madeira basins compared to Amazonas (β = −0.92 and −0.78, pMPE < 0.001 and 0.002, respectively; Table S35, S36), and Ne was lower in Branco (β = −0.34, 95% CI: −0.65 to −0.01, pMPE = 0.04; Tables S33, S34), but basin effects were largely absent for heterozygosity (Table S37, S38).

**Figure 4.**
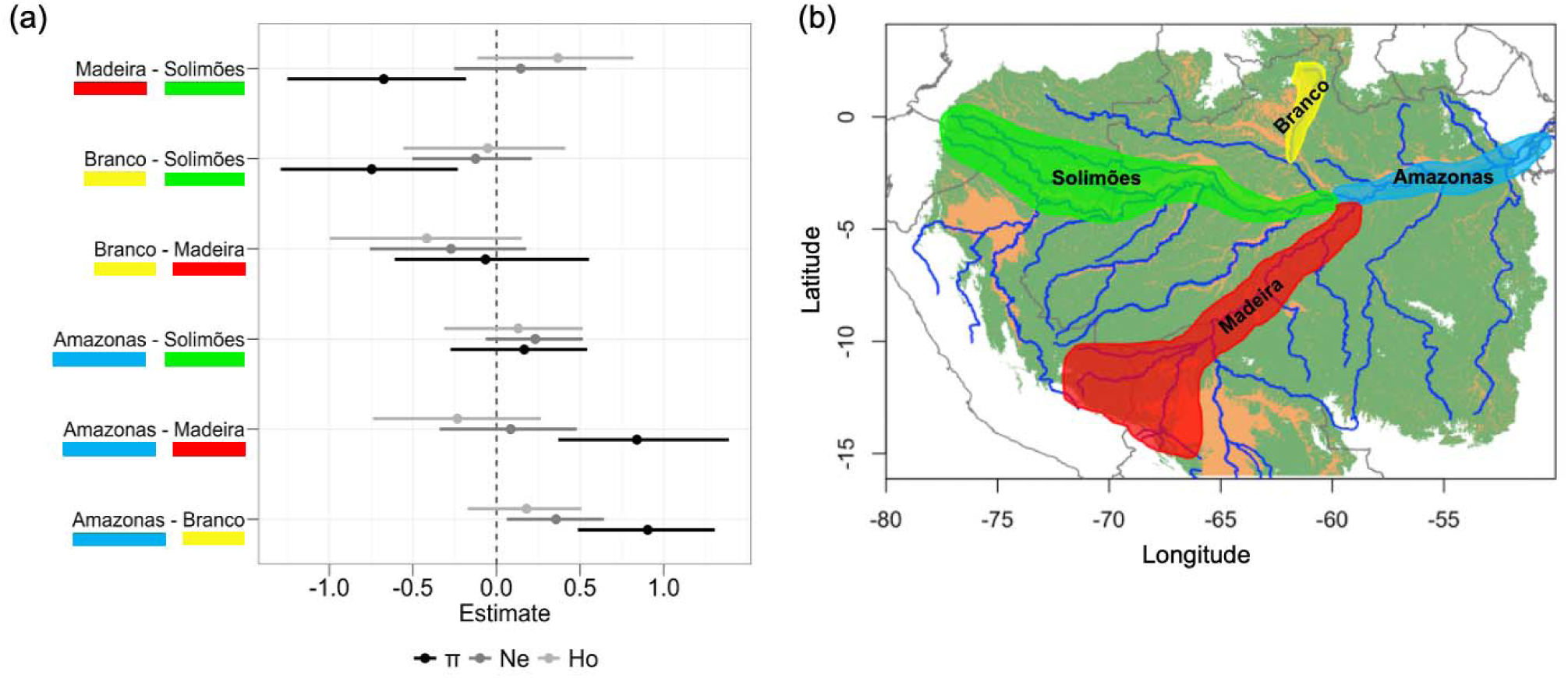
Comparison of genetic diversity (π and H□) and effective population size (Ne) among populations inhabiting major Amazonian sub-basins. (a) Posterior estimates from Bayesian multilevel phylogenetic models comparing genetic diversity and effective population size across sub-basins. (b) Map highlighting the focal sub-basins—Branco, Solimões, Amazonas, and Madeira—used in the comparisons.

## DISCUSSION

### Differential dispersal mediated by habitat specialization shapes spatial genetic variation

Species’ habitat associations provide a direct link between ecological processes and evolutionary change. Because habitat availability strongly influences dispersal (Bowler & Benton, 2005; McPeek & Holt, 1992; Moore et al., 2008), differences in habitat use and movement lead to variation in connectivity and population differentiation. Our results demonstrate that habitat specialization is a major determinant of connectivity in Amazonian river-island birds, shaping patterns of genetic structure and population differentiation. Species associated with ephemeral sandbar scrub habitats have higher dispersal and weaker spatial genetic structure, whereas river-edge forest specialists occupy more stable habitats and show greater genetic differentiation (Figure 3). These differences occur despite the species predominantly sharing similar demographic histories of isolation followed by secondary contact (Table 2), suggesting that habitat specialization affects the degree of population divergence rather than the demographic processes themselves.

The differential contributions of habitat specialization to genetic structure highlight how ecological traits explain distinct dimensions of spatial genetic variation within biological communities. The association between habitat type and genetic structure likely reflects behavioral and ecological responses to seasonal flooding. Sandbar scrub specialists depend on early successional vegetation such as *Tessaria* and *Salix*, which emerge, shift, and disappear across flood cycles (Rosenberg, 1990). As a result, individuals must repeatedly track spatially dynamic habitat patches, imposing stronger mobility than that experienced by species associated with more stable river-edge forest vegetation (Rosenberg, 1990; Rowedder et al., 2021).

Evidence from other avian systems also supports a link between habitat stability and genetic structure. In Amazonian upland forests, vertical habitat use predicts genetic differentiation, with understory species showing stronger population structure than canopy species (Burney & Brumfield, 2009; Smith et al., 2014). This pattern likely reflects differences in resource dynamics. Canopy species track patchy and seasonally variable fruit resources, promoting dispersal, whereas understory insectivores rely on more stable resources and move less, resulting in greater genetic differentiation (Burney & Brumfield, 2009). Similarly, in Central American tropical forests, diet predicts genetic divergence, with plant-dependent species showing lower differentiation than insectivorous or mixed-diet species because they track seasonally changing food resources (Miller et al., 2020). In North American boreal forests, migratory strategy is also associated with genetic structure, as long-distance migrants wintering in relatively stable tropical environments show stronger genetic differentiation than resident or short-distance migrants exposed to more variable high-latitude conditions (Pegan et al., 2025). Our Amazonian river-island system extends these findings by showing that, in highly dynamic floodplains, habitat specialization on shrub or forest vegetation is the main predictor of genetic structure. These habitats differ markedly in their spatial availability across seasonal flood cycles and over the long-term evolution of Amazonian landscapes. Together, these studies suggest that in heterogeneous and seasonal environments, genetic structure is shaped by ecological traits that determine how species respond to spatiotemporal variation in resource availability and habitat stability.

### Temporal connectivity decouples population size and genetic diversity

Besides influencing patterns of spatial genetic variation, habitat association also mediates the relationship among effective population size, gene flow, and genetic diversity. Sandbar scrub populations exhibit smaller effective sizes yet higher nucleotide diversity and observed heterozygosity compared to river-edge forest species (Figure 3c, d). Under equilibrium expectations, reduced population size should lead to a loss of genetic diversity through drift. The opposite pattern observed here instead points to non-equilibrium demographic and genetic dynamics in the Amazonian riverine island system.

In non-equilibrium systems, populations experience repeated changes in size and connectivity instead of remaining stable over time (Excoffier et al., 2009; Marko & Hart, 2011). As a result, effective population size and genetic diversity are not always closely linked because genetic variation reflects both past isolation and subsequent reconnection among populations. This means that genetic diversity can remain high even when contemporary population sizes are small (Alcalá et al., 2013; Alcalá & Vuilleumier, 2014; Hill et al., 2023), supporting the idea that fluctuating environments help maintain genetic and species diversity over time (Yamamichi et al., 2023). Our results are consistent with this prediction. Populations of sandbar specialists species have smaller effective sizes and positive Tajima’s *D* values, indicating recent demographic contractions, but they also show high migration rates among populations (Figure 3). Together, these patterns suggest repeated cycles of isolation and admixture, in which gene flow following periods of fragmentation has the potential to restore genetic variation. As a result, sandbar specialist populations maintain high local genetic diversity despite strong demographic fluctuations, unlike population of species associated with the more stable river-edge forests.

In contrast, populations of river-edge forest species show lower migration rates and lower genetic diversity despite having larger sizes (Figure 3), consistent with longer periods of isolation and the cumulative effects of genetic drift. At the community level, Amazonian river-island bird assemblages show a nested pattern of species-rich and species-poor sites, reflecting repeated cycles of local extinction and recolonization (Borges et al., 2019). Differences in dispersal ability are closely linked to species distributions and extinction histories (Moore et al., 2008). In this case, frequent habitat turnover may reduce long-term effective population sizes through repeated bottlenecks while maintaining genetic diversity by promoting dispersal and gene flow among populations that become temporarily isolated.

Together, these findings demonstrate how non-equilibrium dynamics can decouple population size from genetic diversity in seasonal environments. Consistent with this interpretation, previous studies of Amazonian riverine birds report stronger signals of recent demographic expansion in sandbar scrub specialists than in river-edge forest species (Barbosa et al., 2021; Sawakuchi et al. 2022; Johnson et al., 2023; Luna et al., 2023; Schultz et al., 2024), reinforcing the view that ephemeral habitats promote oscillating population sizes and connectivity. More broadly, similar processes are likely to operate in other fragmented or seasonal systems, where populations persist at small sizes while undergoing repeated cycles of isolation and reconnection (Alcalá & Vuilleumier, 2014; Hill et al., 2023; Fischer & Lindenmayer, 2007). In strongly fluctuating environments, genetic diversity will depend on temporal configuration of population connectivity as much as, or more than, effective population size.

### River sub-basin provides the template for genetic structure

Our spatial genetic analyses revealed a consistent phylogeographic break in the central Amazon (Figure S3), showing that sub-basin intersections shape broad-scale patterns of genetic differentiation. This break matches the regionalization of Amazonian floodplain bird assemblages, which are also structured by river sections and major confluences (Laranjeiras et al., 2024). Despite this geographic division, species occupying similar habitats across different regions showed parallel patterns of divergence and secondary contact. These events represent recent stages along the divergence continuum, spanning ∼774 to 11.7 ka from the Middle Pleistocene to the Holocene, suggesting that dispersal is more strongly constrained by ecological traits than by sub-basin regional processes. This agrees with our previous work showing that riverine birds specialized in sediment-rich systems (e.g., the Amazonas and Branco sub-basins) have clustered divergence times associated with habitat specialization (Luna et al., 2023).

We also found higher nucleotide diversity and larger effective population sizes in the central rivers, including the Amazonas and Solimões, than in neighboring tributaries such as the Madeira and Branco (Figure 4). This pattern is consistent with the interaction between Quaternary climate oscillations and drainage evolution, which created more persistent habitats in western and central Amazonia while promoting environmental instability in the east, contributing to younger eastern Amazonian lineages (Maia-Braga et al., 2026; Silva et al., 2019). These patterns are consistent with Late Quaternary climatic changes that reshaped floodplain dynamics across the basin (Pupim et al., 2019). During glacial–interglacial transitions, periods of high river discharge and low sea level increase erosion, reducing floodplain extent and connectivity for habitat specialists (Pupim et al., 2019; Sawakuchi et al., 2022). At the sub-basin scale, differences in floodplain extent and persistence likely influenced demographic history by changing the availability of suitable habitat. More stable floodplain systems may therefore have maintained larger and more genetically diverse populations. Although our results support a link between habitat dynamics and demographic history, integrating paleoecological reconstructions with denser population genomic sampling across sub-basins will be necessary to better understand these processes.

## Conclusion

Ecological traits, particularly those related to habitat use and dispersal, play a key role in shaping genetic variation and demographic history. Specialization on ephemeral habitats promotes higher dispersal, reduces long-term population structure, and decouples effective population size from genetic diversity. In contrast, specialization on more stable habitats promotes population isolation and greater genetic divergence. As climate change increases environmental variability, many ecosystems—especially in the tropics—are becoming more dynamic and increasingly vulnerable to local extinctions (Lapola et al., 2023; Miles et al., 2004; Pecl et al., 2017). Under these conditions, predicting evolutionary and demographic responses requires integrating ecological traits with models that account for changing landscape connectivity (Vázquez et al., 2017). Static views of landscapes may therefore overlook how shifting habitats shape evolutionary processes through time. By combining habitat associations with comparative population genomics, we show that patterns of genetic divergence and demographic history in seasonal Amazonian landscapes are driven by habitat-mediated dispersal across space and time. More broadly, integrating community ecology with population genetics will be essential for predicting how ongoing environmental change will affect the persistence and diversity of biological communities.

## Acknowledgements

We thank the curators and staff of the Academy of Natural Science of Drexel University, Pennsylvania, USA, Louisiana State University Museum of Natural Sciences, Louisiana, USA, Museu Paraense Emílio Goeldi, Belém, Brazil, and the Instituto Nacional de Pesquisa da Amazônia, Manaus, Brazil for providing tissue samples used in this study. Specimens sequenced in this study were obtained along several decades through collecting permits issued by national environmental agencies in Bolivia, Brazil, and Peru. Financial support to this study was provided by the US Agency for International Development through a Partnerships for Enhanced Engagement in Research grant (Co Ag AID-OAA-A-11-00012), associated with the project "History and diversification of floodplain forest bird communities in Amazonia: Towards an integrated conservation plan". LWL was supported by the fellowship of the Coordenação de Aperfeiçoamento Pessoal de Nível Superior (88882.444617/2019-01), through the Doctoral Graduate Program in Zoology, Universidade Federal do Pará and Museu Paraense Emílio Goeldi. CCR was supported by Brazilian Research Federal and Estate council, FAPEAM and CNPq (research productivity fellowship 314860/2023-1; grants 403708/2024-9; 200531/2025-5). AA is supported by CNPq productivity fellowship (grant 309243/2023-8).

## Notes

### Competing Interest Statement

The authors have declared no competing interest.

