## Supplemental Material for "Habitat specialization structures population divergence and demographic variation in Amazonian floodplain birds"

Table of Content

| **Tables & Figures** | **Content** | **Page** |
| --- | --- | --- |
| **Table S1** | List of species, samples, and localities | 2-18 |
| **Table S2** | Sequence variation summary | 29 |
| **Table S3-S14** | Contribution of variation from each genetic population | 30-34 |
| **Table S15** | Population divergence Weir & Cockerham’s *F*_ST_ | 35 |
| **Table S16** | Genetic diversity for populations grouped by sub-basins | 36 |
| **Table S17-S28** | Estimated demographic parameters under the best-fit model | 37-43 |
| **Table S29-S40** | Results of the Bayesian phylogenetic multilevel model | 44-48 |
| **Figure S1** | Map distribution of samples | 49 |
| **Figure S2** | Cross-validation results from *Construct* analysis comparing spatial and non-spatial models | 50-53 |
| **Figure S3** | Optimal number of genetic populations and genetic-geographic similarity | 54-57 |

**Table S1.** List of bird samples analyzed, including species names, the institution where tissue samples are deposited, voucher sample IDs, the river sub-basin of collection, geographic coordinates (latitude and longitude), and the original reference for the genetic sequence data.

| Species | Institution | Vouch Sample ID | River sub-basin | Latitude | Longitude | Reference |
| --- | --- | --- | --- | --- | --- | --- |
| *Conirostruim bicolor* | INPA | A1043 | Branco | 0.9847 | -61.3413 | Luna et al. 2023 |
| *Conirostruim bicolor* | INPA | A1341 | Japurá | -2.8761 | -64.9193 | Luna et al. 2023 |
| *Conirostruim bicolor* | INPA | A18065 | Japurá | -1.7925 | -68.8189 | Luna et al. 2023 |
| *Conirostruim bicolor* | INPA | A2156 | Branco | 1.7207 | -61.1675 | Luna et al. 2023 |
| *Conirostruim bicolor* | INPA | A23424 | Solimões | -3.1249 | -64.7756 | Luna et al. 2023 |
| *Conirostruim bicolor* | INPA | A23757 | Solimões | -3.8134 | -62.1964 | Luna et al. 2023 |
| *Conirostruim bicolor* | INPA | A23876 | Solimões | -3.3832 | -60.7258 | Luna et al. 2023 |
| *Conirostruim bicolor* | INPA | A23989 | Amazonas | -2.2197 | -54.7548 | Luna et al. 2023 |
| *Conirostruim bicolor* | INPA | A8273 | Branco | 1.5497 | -61.25 | Luna et al. 2023 |
| *Conirostruim bicolor* | INPA | A8279 | Branco | 1.6773 | -61.2001 | Luna et al. 2023 |
| *Conirostruim bicolor* | INPA | B7282 | Solimões | -3.4582 | -72.675 | Luna et al. 2023 |
| *Conirostruim bicolor* | INPA | T1161 | Branco | 1.4313 | -61.2791 | Luna et al. 2023 |
| *Conirostruim bicolor* | INPA | T25522 | Solimões | -3.1086 | -67.9645 | Luna et al. 2023 |
| *Conirostruim bicolor* | INPA | T25524 | Solimões | -3.1086 | -67.9645 | Luna et al. 2023 |
| *Conirostruim bicolor* | INPA | T611 | Branco | 0.7671 | -61.4816 | Luna et al. 2023 |
| *Dendroplex kienerii* | LSU | 20237 | Negro | -2.7501 | -60.75 | Schultz et al., 2023 |
| *Dendroplex kienerii* | LSU | 25477 | Negro | -1.865 | -61.4333 | Schultz et al., 2023 |
| *Dendroplex kienerii* | LSU | 29016 | Solimões | -3.6833 | -73.2 | Schultz et al., 2023 |
| *Dendroplex kienerii* | LSU | 35627 | Amazonas | -2.3761 | -54.6613 | Schultz et al., 2023 |
| *Dendroplex kienerii* | LSU | 35662 | Solimões | -3.3336 | -60.2206 | Schultz et al., 2023 |
| *Dendroplex kienerii* | LSU | 35692 | Solimões | -3.4321 | -68.957 | Schultz et al., 2023 |
| *Dendroplex kienerii* | LSU | 80430 | Solimões | -3.4323 | -68.9571 | Schultz et al., 2023 |
| *Dendroplex kienerii* | LSU | 93459 | Huallaga | -5.4093 | -75.8261 | Schultz et al., 2023 |
| *Dendroplex kienerii* | INPA | A12757 | Tapajós | -4.7498 | -56.6093 | Schultz et al., 2023 |
| *Dendroplex kienerii* | INPA | A2184 | Branco | 1.4572 | -61.2509 | Schultz et al., 2023 |
| *Dendroplex kienerii* | INPA | A23370 | Solimões | -3.0329 | -64.8666 | Schultz et al., 2023 |
| *Dendroplex kienerii* | INPA | A23780 | Solimões | -3.715 | -61.478 | Schultz et al., 2023 |
| *Dendroplex kienerii* | INPA | A23927 | Solimões | -3.6386 | -60.8758 | Schultz et al., 2023 |
| *Dendroplex kienerii* | INPA | A6427 | Negro | -0.4 | -64.8 | Schultz et al., 2023 |
| *Dendroplex kienerii* | INPA | A8321 | Branco | 1.0975 | -61.3272 | Schultz et al., 2023 |
| *Dendroplex kienerii* | INPA | A8341 | Branco | 0.3394 | -61.7508 | Schultz et al., 2023 |
| *Dendroplex kienerii* | MPEG | T16004 | Madeira | -6.7344 | -62.3519 | Schultz et al., 2023 |
| *Dendroplex kienerii* | MPEG | T18603 | Tapajós | -5.7383 | -57.3553 | Schultz et al., 2023 |
| *Dendroplex kienerii* | MPEG | T6618 | Amazonas | -2.0836 | -56.025 | Schultz et al., 2023 |
| *Furnarius minor* | INPA | A1339 | Solimões | -2.7305 | -64.9668 | Schultz et al., 2023 |
| *Furnarius minor* | LSU | B3641 | Solimões | -3.96 | -73.21 | Schultz et al., 2023 |
| *Furnarius minor* | INPA | A23823 | Solimões | -3.666 | -61.5112 | Schultz et al., 2023 |
| *Furnarius minor* | MPEG | MAD489 | Madeira | -4.1247 | -59.3632 | Schultz et al., 2023 |
| *Furnarius minor* | INPA | A23805 | Solimões | -3.666 | -61.5112 | Schultz et al., 2023 |
| *Furnarius minor* | INPA | A23710 | Solimões | -3.8132 | -62.1962 | Schultz et al., 2023 |
| *Furnarius minor* | MPEG | WM083 | Amazonas | -2.0068 | -54.7438 | Schultz et al., 2023 |
| *Furnarius minor* | MPEG | MAD490 | Madeira | -4.1247 | -59.3632 | Schultz et al., 2023 |
| *Furnarius minor* | LSU | B45888 | Amazonas | -2.87 | -58.14 | Schultz et al., 2023 |
| *Furnarius minor* | LSU | B74803 | Ucayali | -10.71 | -73.756 | Schultz et al., 2023 |
| *Serpophaga hypoleuca* | ANSP | 19341 | Napo | -1.0333 | -77.85 | Schultz et al., 2023 |
| *Serpophaga hypoleuca* | LSU | 43033 | Maranõn | -4.7497 | -77.0633 | Schultz et al., 2023 |
| *Serpophaga hypoleuca* | LSU | 43035 | Maranõn | -4.7497 | -77.0633 | Schultz et al., 2023 |
| *Serpophaga hypoleuca* | LSU | 43048 | Maranõn | -4.7497 | -77.0633 | Schultz et al., 2023 |
| *Serpophaga hypoleuca* | LSU | 89189 | Ucayali | -10.6419 | -73.8408 | Schultz et al., 2023 |
| *Serpophaga hypoleuca* | LSU | 89216 | Ucayali | -10.6419 | -73.8408 | Schultz et al., 2023 |
| *Serpophaga hypoleuca* | LSU | 89221 | Ucayali | -10.6419 | -73.8408 | Schultz et al., 2023 |
| *Serpophaga hypoleuca* | INPA | A1342 | Japurá | -2.7306 | -64.9667 | Schultz et al., 2023 |
| *Serpophaga hypoleuca* | INPA | A1480 | Solimões | -3.2251 | -59.9317 | Schultz et al., 2023 |
| *Serpophaga hypoleuca* | INPA | A15959 | Solimões | -3.224 | -59.045 | Schultz et al., 2023 |
| *Serpophaga hypoleuca* | INPA | A23818 | Solimões | -3.666 | -61.5111 | Schultz et al., 2023 |
| *Serpophaga hypoleuca* | INPA | A23821 | Solimões | -3.666 | -61.5111 | Schultz et al., 2023 |
| *Serpophaga hypoleuca* | INPA | A23871 | Solimões | -3.5421 | -60.804 | Schultz et al., 2023 |
| *Serpophaga hypoleuca* | INPA | A23993 | Solimões | -3.5421 | -60.804 | Schultz et al., 2023 |
| *Serpophaga hypoleuca* | INPA | A8302 | Branco | 1.5797 | -61.2472 | Schultz et al., 2023 |
| *Serpophaga hypoleuca* | INPA | A8386 | Branco | -1.0244 | -61.8756 | Schultz et al., 2023 |
| *Cranioleuca vulpecula* | LSU | 3181 | Napo | -3 | -73.3333 | Schultz et al., 2023 |
| *Cranioleuca vulpecula* | LSU | 3637 | Solimões | -3.8756 | -73.2197 | Schultz et al., 2023 |
| *Cranioleuca vulpecula* | LSU | 3643 | Solimões | -3.8756 | -73.2197 | Schultz et al., 2023 |
| *Cranioleuca vulpecula* | LSU | 43020 | Marañon | -4.7497 | -77.0633 | Schultz et al., 2023 |
| *Cranioleuca vulpecula* | LSU | 43025 | Marañon | -4.7497 | -77.0633 | Schultz et al., 2023 |
| *Cranioleuca vulpecula* | LSU | 7253 | Solimões | -3.473 | -72.676 | Schultz et al., 2023 |
| *Cranioleuca vulpecula* | LSU | 7275 | Solimões | -3.473 | -72.676 | Schultz et al., 2023 |
| *Cranioleuca vulpecula* | LSU | 7372 | Solimões | -3.5167 | -72.5167 | Schultz et al., 2023 |
| *Cranioleuca vulpecula* | LSU | 74117 | Ucayali | -10.7469 | -73.7458 | Schultz et al., 2023 |
| *Cranioleuca vulpecula* | LSU | 74802 | Ucayali | -10.71 | -73.76 | Schultz et al., 2023 |
| *Cranioleuca vulpecula* | LSU | 75850 | Ucayali | -10.41 | -73.95 | Schultz et al., 2023 |
| *Cranioleuca vulpecula* | LSU | 89190 | Ucayali | -10.6419 | -73.8408 | Schultz et al., 2023 |
| *Cranioleuca vulpecula* | LSU | 89255 | Ucayali | -10.675 | -73.7969 | Schultz et al., 2023 |
| *Cranioleuca vulpecula* | INPA | A123 | Madeira | -8.8389 | -64.0292 | Schultz et al., 2023 |
| *Cranioleuca vulpecula* | INPA | A124 | Madeira | -8.8389 | -64.0292 | Schultz et al., 2023 |
| *Cranioleuca vulpecula* | INPA | A23717 | Solimões | -3.8133 | -62.1964 | Schultz et al., 2023 |
| *Cranioleuca vulpecula* | INPA | A23720 | Solimões | -3.8133 | -62.1964 | Schultz et al., 2023 |
| *Cranioleuca vulpecula* | INPA | A23801 | Solimões | -3.666 | -61.5111 | Schultz et al., 2023 |
| *Cranioleuca vulpecula* | INPA | A23816 | Solimões | -3.666 | -61.5111 | Schultz et al., 2023 |
| *Cranioleuca vulpecula* | INPA | A23825 | Solimões | -3.666 | -61.5111 | Schultz et al., 2023 |
| *Cranioleuca vulpecula* | INPA | A23838 | Solimões | -3.666 | -61.5111 | Schultz et al., 2023 |
| *Cranioleuca vulpecula* | LSU | 25424 | Solimões | -3.25375 | -59.9719 | Schultz et al., 2023 |
| *Cranioleuca vulpecula* | LSU | 74799 | Ucayali | -10.6853 | -73.7968 | Schultz et al., 2023 |
| *Cranioleuca vulpecula* | LSU | 79790 | Madre de Dios | -10.992 | -66.083 | Schultz et al., 2023 |
| *Cranioleuca vulpecula* | LSU | 93331 | Madre de Dios | -5.399 | -75.801 | Schultz et al., 2023 |
| *Cranioleuca vulpecula* | MPEG | T16311 | Madeira | -4.1283 | -59.3653 | Schultz et al., 2023 |
| *Mazaria propinqua* | LSU | 43070 | Marañon | -4.7497 | -77.0633 | Luna et al. 2023 |
| *Mazaria propinqua* | LSU | 43083 | Marañon | -4.7497 | -77.0633 | Luna et al. 2023 |
| *Mazaria propinqua* | LSU | 7289 | Solimões | -4.4585 | -73.4996 | Luna et al. 2023 |
| *Mazaria propinqua* | LSU | 7331 | Solimões | -3.5167 | -72.5167 | Luna et al. 2023 |
| *Mazaria propinqua* | LSU | 79791 | Madre de Dios | -10.9919 | -66.0831 | Luna et al. 2023 |
| *Mazaria propinqua* | LSU | 79801 | Madre de Dios | -11.1819 | -65.7628 | Luna et al. 2023 |
| *Mazaria propinqua* | LSU | 89182 | Ucayali | -10.6419 | -73.8408 | Luna et al. 2023 |
| *Mazaria propinqua* | LSU | 89264 | Ucayali | -10.675 | -73.7969 | Luna et al. 2023 |
| *Mazaria propinqua* | INPA | A1025 | Branco | 1.5483 | -61.2553 | Luna et al. 2023 |
| *Mazaria propinqua* | INPA | A18287 | Japurá | -1.7167 | -69.1167 | Luna et al. 2023 |
| *Mazaria propinqua* | INPA | A2158 | Branco | 1.5576 | -61.2489 | Luna et al. 2023 |
| *Mazaria propinqua* | INPA | A23479 | Solimões | -3.0961 | -64.757 | Luna et al. 2023 |
| *Mazaria propinqua* | INPA | A23707 | Solimões | -3.8132 | -62.1962 | Luna et al. 2023 |
| *Mazaria propinqua* | INPA | A23708 | Solimões | -3.8132 | -62.1962 | Luna et al. 2023 |
| *Mazaria propinqua* | INPA | A23872 | Solimões | -3.5421 | -60.804 | Luna et al. 2023 |
| *Mazaria propinqua* | INPA | A23886 | Solimões | -3.5421 | -60.804 | Luna et al. 2023 |
| *Mazaria propinqua* | INPA | A8309 | Branco | 1.1665 | -61.3341 | Luna et al. 2023 |
| *Mazaria propinqua* | INPA | A8345 | Branco | 0.2664 | -61.7653 | Luna et al. 2023 |
| *Mazaria propinqua* | MPEG | T1027 | Branco | 1.482 | -61.253 | Luna et al. 2023 |
| *Mazaria propinqua* | MPEG | T760 | Branco | -0.6155 | -61.8133 | Luna et al. 2023 |
| *Mazaria propinqua* | LSU | 763 | Branco | -0.6155 | -61.8133 | Luna et al. 2023 |
| *Mazaria propinqua* | LSU | 765 | Branco | -0.6138 | -61.8122 | Luna et al. 2023 |
| *Mazaria propinqua* | MPEG | T767 | Branco | 1.482 | -61.253 | Luna et al. 2023 |
| *Mazaria propinqua* | LSU | 43070 | Marañon | -4.7497 | -77.0633 | Luna et al. 2023 |
| *Mazaria propinqua* | LSU | 43083 | Marañon | -4.7497 | -77.0633 | Luna et al. 2023 |
| *Mazaria propinqua* | LSU | 7289 | Solimões | -4.4585 | -73.4996 | Luna et al. 2023 |
| *Mazaria propinqua* | LSU | 7331 | Solimões | -3.5167 | -72.5167 | Luna et al. 2023 |
| *Stigmatura napensis* | LSU | 17725 | Napo | -1.0333 | -77.85 | Luna et al. 2023 |
| *Stigmatura napensis* | LSU | 19369 | Napo | -1.0333 | -77.85 | Luna et al. 2023 |
| *Stigmatura napensis* | LSU | 3638 | Solimões | -3.8756 | -73.2197 | Luna et al. 2023 |
| *Stigmatura napensis* | LSU | 3639 | Solimões | -3.8756 | -73.2197 | Luna et al. 2023 |
| *Stigmatura napensis* | LSU | 43079 | Marañon | -4.7497 | -77.13 | Luna et al. 2023 |
| *Stigmatura napensis* | LSU | 7240 | Solimões | -3.473 | -72.676 | Luna et al. 2023 |
| *Stigmatura napensis* | LSU | 89188 | Ucayali | -10.6419 | -73.8408 | Luna et al. 2023 |
| *Stigmatura napensis* | LSU | 89217 | Ucayali | -10.6419 | -73.8408 | Luna et al. 2023 |
| *Stigmatura napensis* | LSU | 89218 | Ucayali | -10.6419 | -73.8408 | Luna et al. 2023 |
| *Stigmatura napensis* | INPA | A027 | Solimões | -2.6806 | -66.8611 | Luna et al. 2023 |
| *Stigmatura napensis* | INPA | A028 | Solimões | -2.6806 | -66.8611 | Luna et al. 2023 |
| *Stigmatura napensis* | INPA | A1033 | Branco | 1.49933 | -61.2527 | Luna et al. 2023 |
| *Stigmatura napensis* | INPA | A14198 | Branco | -1.0047 | -61.8812 | Luna et al. 2023 |
| *Stigmatura napensis* | INPA | A15092 | Tapajós | -3.3 | -55.2833 | Luna et al. 2023 |
| *Stigmatura napensis* | INPA | A15126 | Tapajós | -3.3167 | -55.3333 | Luna et al. 2023 |
| *Stigmatura napensis* | INPA | A15923 | Branco | -0.0496 | -61.8074 | Luna et al. 2023 |
| *Stigmatura napensis* | INPA | A15960 | Solimões | -3.24 | -59.0603 | Luna et al. 2023 |
| *Stigmatura napensis* | INPA | A2166 | Branco | 1.4673 | -61.2536 | Luna et al. 2023 |
| *Stigmatura napensis* | INPA | A23482 | Solimões | -3.0961 | -64.757 | Luna et al. 2023 |
| *Stigmatura napensis* | INPA | A23483 | Solimões | -3.0961 | -64.757 | Luna et al. 2023 |
| *Stigmatura napensis* | INPA | A23703 | Solimões | -3.8133 | -62.1964 | Luna et al. 2023 |
| *Stigmatura napensis* | INPA | A23711 | Solimões | -3.8133 | -62.1964 | Luna et al. 2023 |
| *Stigmatura napensis* | INPA | A23892 | Solimões | -3.5421 | -60.804 | Luna et al. 2023 |
| *Stigmatura napensis* | INPA | A23895 | Solimões | -3.5421 | -60.804 | Luna et al. 2023 |
| *Stigmatura napensis* | INPA | A8280 | Branco | 1.5261 | -61.2467 | Luna et al. 2023 |
| *Stigmatura napensis* | INPA | A8330 | Branco | 0.5075 | -61.6647 | Luna et al. 2023 |
| *Stigmatura napensis* | INPA | A8385 | Branco | -1.0244 | -61.8756 | Luna et al. 2023 |
| *Stigmatura napensis* | MPEG | T16303 | Madeira | -4.3825 | -59.7517 | Luna et al. 2023 |
| *Stigmatura napensis* | MPEG | T16335 | Madeira | -4.1283 | -59.3653 | Luna et al. 2023 |
| *Myrmochanes hemileucus* | LSU | 3182 | Napo | -3 | -73.3333 | Schultz et al., 2023 |
| *Myrmochanes hemileucus* | LSU | 3268 | Napo | -3 | -73.3667 | Schultz et al., 2023 |
| *Myrmochanes hemileucus* | LSU | 3644 | Solimões | -3.8756 | -73.2197 | Schultz et al., 2023 |
| *Myrmochanes hemileucus* | LSU | 43034 | Marañon | -4.7497 | -77.0633 | Schultz et al., 2023 |
| *Myrmochanes hemileucus* | LSU | 43066 | Marañon | -4.7497 | -77.0633 | Schultz et al., 2023 |
| *Myrmochanes hemileucus* | LSU | 7245 | Solimões | -3.473 | -72.676 | Schultz et al., 2023 |
| *Myrmochanes hemileucus* | LSU | 74113 | Ucayali | -10.7469 | -73.7458 | Schultz et al., 2023 |
| *Myrmochanes hemileucus* | LSU | 74773 | Ucayali | -10.71 | -73.76 | Schultz et al., 2023 |
| *Myrmochanes hemileucus* | LSU | 89240 | Ucayali | -10.675 | -73.7969 | Schultz et al., 2023 |
| *Myrmochanes hemileucus* | INPA | A11135 | Solimões | -3.455 | -60.7733 | Schultz et al., 2023 |
| *Myrmochanes hemileucus* | INPA | A1340 | Japurá | -2.7306 | -64.9667 | Schultz et al., 2023 |
| *Myrmochanes hemileucus* | INPA | A23393 | Solimões | -3.1245 | -64.7751 | Schultz et al., 2023 |
| *Myrmochanes hemileucus* | INPA | A23394 | Solimões | -3.1245 | -64.7751 | Schultz et al., 2023 |
| *Myrmochanes hemileucus* | INPA | A23709 | Solimões | -3.8133 | -62.1965 | Schultz et al., 2023 |
| *Myrmochanes hemileucus* | INPA | A23726 | Solimões | -3.8133 | -62.1965 | Schultz et al., 2023 |
| *Myrmochanes hemileucus* | INPA | A23804 | Solimões | -3.666 | -61.5112 | Schultz et al., 2023 |
| *Myrmochanes hemileucus* | MPEG | T16307 | Madeira | -4.1283 | -59.3653 | Schultz et al., 2023 |
| *Myrmochanes hemileucus* | MPEG | T16339 | Madeira | -4.1283 | -59.3653 | Schultz et al., 2023 |
| *Myrmochanes hemileucus* | MPEG | T22543 | Solimões | -3.1478 | -67.9597 | Schultz et al., 2023 |
| *Myrmochanes hemileucus* | MPEG | T22561 | Solimões | -2.4692 | -66.0858 | Schultz et al., 2023 |
| *Myrmochanes hemileucus* | MPEG | T22712 | Solimões | -3.8433 | -62.2414 | Schultz et al., 2023 |
| *Myrmotherula klagesi* | INPA | A048 | Solimões | -3.9028 | -62.8222 | Schultz et al., 2023 |
| *Myrmotherula klagesi* | INPA | A1044 | Branco | 0.9578 | -61.355 | Schultz et al., 2023 |
| *Myrmotherula klagesi* | INPA | A10501 | Negro | -1.8438 | -61.3801 | Schultz et al., 2023 |
| *Myrmotherula klagesi* | INPA | A10588 | Negro | -1.8757 | -61.3671 | Schultz et al., 2023 |
| *Myrmotherula klagesi* | INPA | A15925 | Branco | -0.1083 | -61.8125 | Schultz et al., 2023 |
| *Myrmotherula klagesi* | INPA | A4907 | Negro | -1.8742 | -61.3692 | Schultz et al., 2023 |
| *Myrmotherula klagesi* | INPA | A8276 | Branco | 1.5311 | -61.2456 | Schultz et al., 2023 |
| *Myrmotherula klagesi* | INPA | A8413 | Branco | -0.9881 | -61.8908 | Schultz et al., 2023 |
| *Myrmotherula klagesi* | INPA | A967 | Solimões | -3.7811 | -64.0254 | Schultz et al., 2023 |
| *Myrmotherula klagesi* | INPA | A968 | Solimões | -3.7811 | -64.0254 | Schultz et al., 2023 |
| *Myrmotherula klagesi* | LSU | 20250 | Negro | -2.75 | -60.75 | Schultz et al., 2023 |
| *Myrmotherula klagesi* | LSU | 25511 | Negro | -1.9756 | -61.2686 | Schultz et al., 2023 |
| *Myrmotherula klagesi* | LSU | 25560 | Amazonas | -2.8342 | -58.1411 | Schultz et al., 2023 |
| *Myrmotherula klagesi* | LSU | 25562 | Amazonas | -2.8342 | -58.1411 | Schultz et al., 2023 |
| *Myrmotherula klagesi* | LSU | 81357 | Madeira | -3.5922 | -58.9383 | Schultz et al., 2023 |
| *Myrmotherula klagesi* | LSU | 81365 | Madeira | -3.5922 | -58.9383 | Schultz et al., 2023 |
| *Myrmotherula klagesi* | MPEG | T16396 | Madeira | -3.5919 | -58.9431 | Schultz et al., 2023 |
| *Myrmotherula klagesi* | MPEG | T22645 | Solimões | -4.0136 | -62.95 | Schultz et al., 2023 |
| *Myrmotherula klagesi* | MPEG | T22836 | Amazonas | -3.1586 | -58.3703 | Schultz et al., 2023 |
| *Myrmotherula assimilis* | MPEG | ETA046 | Solimões | -4.3244 | -69.8862 | Thom et al. 2020 |
| *Myrmotherula assimilis* | MPEG | ETA051 | Solimões | -4.3244 | -69.8862 | Thom et al. 2020 |
| *Myrmotherula assimilis* | MPEG | ETA054 | Solimões | -4.3244 | -69.8862 | Thom et al. 2020 |
| *Myrmotherula assimilis* | MPEG | ETA081 | Solimões | -3.148 | -67.9598 | Thom et al. 2020 |
| *Myrmotherula assimilis* | MPEG | ETA097 | Solimões | -3.148 | -67.9598 | Thom et al. 2020 |
| *Myrmotherula assimilis* | MPEG | ETA098 | Solimões | -3.148 | -67.9598 | Thom et al. 2020 |
| *Myrmotherula assimilis* | MPEG | ETA212 | Solimões | -3.3794 | -64.6411 | Thom et al. 2020 |
| *Myrmotherula assimilis* | MPEG | ETA254 | Solimões | -4.0137 | -62.9501 | Thom et al. 2020 |
| *Myrmotherula assimilis* | MPEG | ETA269 | Solimões | -4.0137 | -62.9501 | Thom et al. 2020 |
| *Myrmotherula assimilis* | MPEG | ETA337 | Solimões | -3.2878 | -60.215 | Thom et al. 2020 |
| *Myrmotherula assimilis* | MPEG | ETA338 | Solimões | -3.2878 | -60.215 | Thom et al. 2020 |
| *Myrmotherula assimilis* | MPEG | ETA339 | Solimões | -3.2878 | -60.215 | Thom et al. 2020 |
| *Myrmotherula assimilis* | MPEG | ETA353 | Solimões | -3.2878 | -60.215 | Thom et al. 2020 |
| *Myrmotherula assimilis* | MPEG | ETA399 | Madeira | -4.3755 | -59.5971 | Thom et al. 2020 |
| *Myrmotherula assimilis* | MPEG | ETA400 | Madeira | -4.3755 | -59.5971 | Thom et al. 2020 |
| *Myrmotherula assimilis* | MPEG | ETA401 | Madeira | -4.3755 | -59.5971 | Thom et al. 2020 |
| *Myrmotherula assimilis* | MPEG | ETA402 | Madeira | -4.3755 | -59.5971 | Thom et al. 2020 |
| *Myrmotherula assimilis* | MPEG | ETA403 | Madeira | -4.3755 | -59.5971 | Thom et al. 2020 |
| *Myrmotherula assimilis* | MPEG | ETA440 | Madeira | -3.5715 | -58.9228 | Thom et al. 2020 |
| *Myrmotherula assimilis* | MPEG | ETA441 | Madeira | -3.5715 | -58.9228 | Thom et al. 2020 |
| *Myrmotherula assimilis* | MPEG | JUR011 | Amazonas | -2.0836 | -56.0253 | Thom et al. 2020 |
| *Myrmotherula assimilis* | MPEG | 56696 | Amazonas | -2.0836 | -56.0253 | Thom et al. 2020 |
| *Myrmotherula assimilis* | INPA | A15788 | Negro | -1.855 | -61.4383 | Thom et al. 2020 |
| *Myrmotherula assimilis* | INPA | A8418 | Negro | -1.0986 | -61.9431 | Thom et al. 2020 |
| *Myrmotherula assimilis* | MPEG | AMA427 | Solimões | -4.4867 | -71.5506 | Thom et al. 2020 |
| *Myrmotherula assimilis* | MPEG | AMA475 | Solimões | -4.4867 | -71.5506 | Thom et al. 2020 |
| *Myrmotherula assimilis* | MPEG | ETA442 | Madeira | -3.5715 | -58.9228 | Thom et al. 2020 |
| *Myrmotherula assimilis* | MPEG | ETA490 | Amazonas | -3.1588 | -58.3703 | Thom et al. 2020 |
| *Myrmotherula assimilis* | MPEG | ETA491 | Amazonas | -3.1588 | -58.3703 | Thom et al. 2020 |
| *Myrmotherula assimilis* | MPEG | ETA509 | Amazonas | -3.1845 | -58.2645 | Thom et al. 2020 |
| *Myrmotherula assimilis* | MPEG | ETA559 | Amazonas | -2.5799 | -56.6792 | Thom et al. 2020 |
| *Myrmotherula assimilis* | MPEG | ETA560 | Amazonas | -2.5799 | -56.6792 | Thom et al. 2020 |
| *Myrmotherula assimilis* | MPEG | JUT269 | Solimões | -3.225 | -67.4361 | Thom et al. 2020 |
| *Myrmotherula assimilis* | MPEG | MAD057 | Madeira | -8.2239 | -63.1818 | Thom et al. 2020 |
| *Myrmotherula assimilis* | MPEG | MAD060 | Madeira | -8.2239 | -63.1818 | Thom et al. 2020 |
| *Myrmotherula assimilis* | MPEG | MAD303 | Madeira | -5.6129 | -61.1216 | Thom et al. 2020 |
| *Myrmotherula assimilis* | MPEG | MAD337 | Madeira | -5.6129 | -61.1216 | Thom et al. 2020 |
| *Myrmotherula assimilis* | MPEG | MSF408 | Madeira | -8.9367 | -63.3514 | Thom et al. 2020 |
| *Thamnophilus cryptoleucus* | MPEG | ETA 003 | Solimões | -4.3243 | -69.8861 | Thom et al. 2020 |
| *Thamnophilus cryptoleucus* | MPEG | ETA 005 | Solimões | -4.3243 | -69.8861 | Thom et al. 2020 |
| *Thamnophilus cryptoleucus* | MPEG | ETA 007 | Solimões | -4.3243 | -69.8861 | Thom et al. 2020 |
| *Thamnophilus cryptoleucus* | MPEG | ETA 008 | Solimões | -4.3243 | -69.8861 | Thom et al. 2020 |
| *Thamnophilus cryptoleucus* | MPEG | ETA 174 | Solimões | -2.4828 | -66.0106 | Thom et al. 2020 |
| *Thamnophilus cryptoleucus* | MPEG | ETA 176 | Solimões | -2.4828 | -66.0106 | Thom et al. 2020 |
| *Thamnophilus cryptoleucus* | MPEG | ETA 240 | Solimões | -4.0137 | -62.9501 | Thom et al. 2020 |
| *Thamnophilus cryptoleucus* | MPEG | ETA 242 | Solimões | -4.0137 | -62.9501 | Thom et al. 2020 |
| *Thamnophilus cryptoleucus* | MPEG | ETA 243 | Solimões | -4.0137 | -62.9501 | Thom et al. 2020 |
| *Thamnophilus cryptoleucus* | MPEG | ETA 331 | Solimões | -3.2877 | -60.215 | Thom et al. 2020 |
| *Thamnophilus cryptoleucus* | MPEG | ETA 332 | Solimões | -3.2877 | -60.215 | Thom et al. 2020 |
| *Thamnophilus cryptoleucus* | MPEG | ETA 335 | Solimões | -3.2877 | -60.215 | Thom et al. 2020 |
| *Thamnophilus cryptoleucus* | MPEG | ETA 062 | Solimões | -3.1479 | -67.9597 | Thom et al. 2020 |
| *Thamnophilus cryptoleucus* | MPEG | ETA 063 | Solimões | -3.1479 | -67.9597 | Thom et al. 2020 |
| *Thamnophilus cryptoleucus* | MPEG | ETA 175 | Solimões | -2.4828 | -66.0106 | Thom et al. 2020 |
| *Thamnophilus cryptoleucus* | MPEG | ETA 201 | Solimões | -3.3794 | -64.641 | Thom et al. 2020 |
| *Thamnophilus cryptoleucus* | MPEG | ETA 202 | Solimões | -3.3794 | -64.641 | Thom et al. 2020 |
| *Thamnophilus cryptoleucus* | MPEG | ETA 289 | Solimões | -3.8434 | -62.2415 | Thom et al. 2020 |
| *Thamnophilus cryptoleucus* | MPEG | ETA 290 | Solimões | -3.8434 | -62.2415 | Thom et al. 2020 |
| *Thamnophilus cryptoleucus* | MPEG | ETA 418 | Madeira | -3.5714 | -58.9228 | Thom et al. 2020 |
| *Thamnophilus cryptoleucus* | MPEG | ETA 419 | Madeira | -3.5714 | -58.9228 | Thom et al. 2020 |
| *Thamnophilus cryptoleucus* | MPEG | ETA 420 | Madeira | -3.5714 | -58.9228 | Thom et al. 2020 |
| *Myrmoborus lugubris* | INPA | A10505 | Negro | -1.8438 | -61.38 | Thom et al. 2020 |
| *Myrmoborus lugubris* | INPA | A10508 | Negro | -1.8756 | -61.3671 | Thom et al. 2020 |
| *Myrmoborus lugubris* | INPA | A15803 | Negro | -1.8102 | -61.395 | Thom et al. 2020 |
| *Myrmoborus lugubris* | INPA | A15805 | Negro | -1.8102 | -61.395 | Thom et al. 2020 |
| *Myrmoborus lugubris* | INPA | A2191 | Branco | 1.2402 | -61.3141 | Thom et al. 2020 |
| *Myrmoborus lugubris* | INPA | A3095 | Negro | -1.8657 | -61.3724 | Thom et al. 2020 |
| *Myrmoborus lugubris* | INPA | A3103 | Negro | -1.8657 | -61.3724 | Thom et al. 2020 |
| *Myrmoborus lugubris* | INPA | A8376 | Branco | -0.53 | -61.7991 | Thom et al. 2020 |
| *Myrmoborus lugubris* | INPA | A8379 | Branco | -0.53 | -61.7991 | Thom et al. 2020 |
| *Myrmoborus lugubris* | INPA | A936 | Solimões | -3.781 | -64.0253 | Thom et al. 2020 |
| *Myrmoborus lugubris* | MPEG | ETA 001 | Solimões | -4.3243 | -69.8861 | Thom et al. 2020 |
| *Myrmoborus lugubris* | MPEG | ETA 002 | Solimões | -4.3243 | -69.8861 | Thom et al. 2020 |
| *Myrmoborus lugubris* | MPEG | ETA 004 | Solimões | -4.3243 | -69.8861 | Thom et al. 2020 |
| *Myrmoborus lugubris* | MPEG | ETA 009 | Solimões | -4.3243 | -69.8861 | Thom et al. 2020 |
| *Myrmoborus lugubris* | MPEG | ETA 052 | Solimões | -4.3243 | -69.8861 | Thom et al. 2020 |
| *Myrmoborus lugubris* | MPEG | ETA 053 | Solimões | -4.3243 | -69.8861 | Thom et al. 2020 |
| *Myrmoborus lugubris* | MPEG | ETA 073 | Solimões | -3.1479 | -67.9597 | Thom et al. 2020 |
| *Myrmoborus lugubris* | MPEG | ETA 074 | Solimões | -3.1479 | -67.9597 | Thom et al. 2020 |
| *Myrmoborus lugubris* | MPEG | ETA 093 | Solimões | -3.1479 | -67.9597 | Thom et al. 2020 |
| *Myrmoborus lugubris* | MPEG | ETA 094 | Solimões | -3.1479 | -67.9597 | Thom et al. 2020 |
| *Myrmoborus lugubris* | MPEG | ETA 096 | Solimões | -3.1479 | -67.9597 | Thom et al. 2020 |
| *Myrmoborus lugubris* | MPEG | ETA 114 | Solimões | -3.1479 | -67.9597 | Thom et al. 2020 |
| *Myrmoborus lugubris* | MPEG | ETA 115 | Solimões | -3.1479 | -67.9597 | Thom et al. 2020 |
| *Myrmoborus lugubris* | MPEG | ETA 162 | Solimões | -2.4828 | -66.0106 | Thom et al. 2020 |
| *Myrmoborus lugubris* | MPEG | ETA 163 | Solimões | -2.4828 | -66.0106 | Thom et al. 2020 |
| *Myrmoborus lugubris* | MPEG | ETA 164 | Solimões | -2.4828 | -66.0106 | Thom et al. 2020 |
| *Myrmoborus lugubris* | MPEG | ETA 165 | Solimões | -2.4828 | -66.0106 | Thom et al. 2020 |
| *Myrmoborus lugubris* | MPEG | ETA 166 | Solimões | -2.4828 | -66.0106 | Thom et al. 2020 |
| *Myrmoborus lugubris* | MPEG | ETA 194 | Solimões | -2.4828 | -66.0106 | Thom et al. 2020 |
| *Myrmoborus lugubris* | MPEG | ETA 196 | Solimões | -2.4828 | -66.0106 | Thom et al. 2020 |
| *Myrmoborus lugubris* | MPEG | ETA 233 | Solimões | -3.2658 | -64.7071 | Thom et al. 2020 |
| *Myrmoborus lugubris* | MPEG | ETA 234 | Solimões | -3.2658 | -64.7071 | Thom et al. 2020 |
| *Myrmoborus lugubris* | MPEG | ETA 235 | Solimões | -3.2658 | -64.7071 | Thom et al. 2020 |
| *Myrmoborus lugubris* | MPEG | ETA 236 | Solimões | -3.2658 | -64.7071 | Thom et al. 2020 |
| *Myrmoborus lugubris* | MPEG | ETA 237 | Solimões | -3.2658 | -64.7071 | Thom et al. 2020 |
| *Myrmoborus lugubris* | MPEG | ETA 238 | Solimões | -3.2658 | -64.7071 | Thom et al. 2020 |
| *Myrmoborus lugubris* | MPEG | ETA 246 | Solimões | -4.0137 | -62.9501 | Thom et al. 2020 |
| *Myrmoborus lugubris* | MPEG | ETA 256 | Solimões | -4.0137 | -62.9501 | Thom et al. 2020 |
| *Myrmoborus lugubris* | MPEG | ETA 257 | Solimões | -4.0137 | -62.9501 | Thom et al. 2020 |
| *Myrmoborus lugubris* | MPEG | ETA 258 | Solimões | -4.0137 | -62.9501 | Thom et al. 2020 |
| *Myrmoborus lugubris* | MPEG | ETA 368 | Madeira | -4.3754 | -59.597 | Thom et al. 2020 |
| *Myrmoborus lugubris* | MPEG | ETA 369 | Madeira | -4.3754 | -59.597 | Thom et al. 2020 |
| *Myrmoborus lugubris* | MPEG | ETA 370 | Madeira | -4.3754 | -59.597 | Thom et al. 2020 |
| *Myrmoborus lugubris* | MPEG | ETA 371 | Madeira | -4.3754 | -59.597 | Thom et al. 2020 |
| *Myrmoborus lugubris* | MPEG | ETA 372 | Madeira | -4.3754 | -59.597 | Thom et al. 2020 |
| *Myrmoborus lugubris* | MPEG | ETA 373 | Madeira | -4.3754 | -59.597 | Thom et al. 2020 |
| *Myrmoborus lugubris* | MPEG | ETA 385 | Madeira | -4.3754 | -59.597 | Thom et al. 2020 |
| *Myrmoborus lugubris* | MPEG | ETA 429 | Madeira | -3.5714 | -58.9228 | Thom et al. 2020 |
| *Myrmoborus lugubris* | MPEG | ETA 430 | Madeira | -3.5714 | -58.9228 | Thom et al. 2020 |
| *Myrmoborus lugubris* | MPEG | ETA 431 | Madeira | -3.5714 | -58.9228 | Thom et al. 2020 |
| *Myrmoborus lugubris* | MPEG | ETA 432 | Madeira | -3.5714 | -58.9228 | Thom et al. 2020 |
| *Myrmoborus lugubris* | MPEG | ETA 433 | Madeira | -3.5714 | -58.9228 | Thom et al. 2020 |
| *Myrmoborus lugubris* | MPEG | ETA 434 | Madeira | -3.5714 | -58.9228 | Thom et al. 2020 |
| *Myrmoborus lugubris* | MPEG | ETA 465 | Amazonas | -3.1587 | -58.3702 | Thom et al. 2020 |
| *Myrmoborus lugubris* | MPEG | ETA 497 | Amazonas | -3.1845 | -58.2644 | Thom et al. 2020 |
| *Myrmoborus lugubris* | MPEG | ETA 498 | Amazonas | -3.1845 | -58.2644 | Thom et al. 2020 |
| *Myrmoborus lugubris* | MPEG | ETA 499 | Amazonas | -3.1845 | -58.2644 | Thom et al. 2020 |
| *Myrmoborus lugubris* | MPEG | ETA 500 | Amazonas | -3.1845 | -58.2644 | Thom et al. 2020 |
| *Myrmoborus lugubris* | MPEG | ETA 501 | Amazonas | -3.1845 | -58.2644 | Thom et al. 2020 |
| *Myrmoborus lugubris* | MPEG | ETA 502 | Amazonas | -3.1845 | -58.2644 | Thom et al. 2020 |
| *Myrmoborus lugubris* | MPEG | ETA 508 | Amazonas | -3.1845 | -58.2644 | Thom et al. 2020 |
| *Myrmoborus lugubris* | MPEG | ETA 512 | Amazonas | -2.5798 | -56.6791 | Thom et al. 2020 |
| *Myrmoborus lugubris* | MPEG | ETA 513 | Amazonas | -2.5798 | -56.6791 | Thom et al. 2020 |
| *Myrmoborus lugubris* | MPEG | ETA 514 | Amazonas | -2.5798 | -56.6791 | Thom et al. 2020 |
| *Myrmoborus lugubris* | MPEG | ETA 515 | Amazonas | -2.5798 | -56.6791 | Thom et al. 2020 |
| *Myrmoborus lugubris* | MPEG | ETA 516 | Amazonas | -2.5798 | -56.6791 | Thom et al. 2020 |
| *Myrmoborus lugubris* | MPEG | ETA 534 | Amazonas | -2.5798 | -56.6791 | Thom et al. 2020 |
| *Myrmoborus lugubris* | MPEG | ETA 563 | Amazonas | -2.3881 | -54.3616 | Thom et al. 2020 |
| *Myrmoborus lugubris* | MPEG | ETA 564 | Amazonas | -2.3881 | -54.3616 | Thom et al. 2020 |
| *Myrmoborus lugubris* | MPEG | ETA 565 | Amazonas | -2.3881 | -54.3616 | Thom et al. 2020 |
| *Myrmoborus lugubris* | MPEG | ETA 566 | Amazonas | -2.3881 | -54.3616 | Thom et al. 2020 |
| *Myrmoborus lugubris* | MPEG | ETA 567 | Amazonas | -2.3881 | -54.3616 | Thom et al. 2020 |
| *Myrmoborus lugubris* | MPEG | ETA 597 | Amazonas | -1.5375 | -52.5238 | Thom et al. 2020 |
| *Myrmoborus lugubris* | MPEG | ETA 598 | Amazonas | -1.5375 | -52.5238 | Thom et al. 2020 |
| *Myrmoborus lugubris* | MPEG | ETA 599 | Amazonas | -1.5375 | -52.5238 | Thom et al. 2020 |
| *Myrmoborus lugubris* | MPEG | ETA 600 | Amazonas | -1.5375 | -52.5238 | Thom et al. 2020 |
| *Myrmoborus lugubris* | MPEG | ETA 601 | Amazonas | -1.5375 | -52.5238 | Thom et al. 2020 |
| *Myrmoborus lugubris* | MPEG | ETA 645 | Amazonas | -1.5375 | -52.5238 | Thom et al. 2020 |
| *Myrmoborus lugubris* | MPEG | ETA 646 | Amazonas | -1.5375 | -52.5238 | Thom et al. 2020 |
| *Myrmoborus lugubris* | INPA | A2193 | Branco | 1.2402 | -61.3141 | Thom et al. 2020 |

**Table S2.** Genetic summary for each bird species with the number of samples (N), number of assembled loci, informative sites, total SNPs for each database, one biallelic SNP per locus with a minimum frequency of 0.01 called from the total database*.*

| Species | N | Loci^1^ | | Informative site | Total SNPs | SNPs one per locus^1^ |
| --- | --- | --- | --- | --- | --- | --- |
| *Conirostrum bicolor* | 15 | 2,339 | 7,536 | | 7,679 | 708 |
| *Dendroplex kienerii* | 19 | 2,349 | 18,019 | | 10,355 | 1,409 |
| *Mazaria proprinqua* | 28 | 2,347 | 11,244 | | 3,732 | 1,107 |
| *Cranioleuca vulpecula* | 26 | 2,362 | 7,577 | | 4,091 | 1,274 |
| *Furnarius minor* | 10 | 2,337 | 14,377 | | 7,914 | 1,108 |
| *Serpophaga hypoleuca* | 16 | 2,346 | 10,345 | | 10,769 | 1,209 |
| *Stigmatura napensis* | 29 | 2,364 | 15,836 | | 8,518 | 1,163 |
| *Myrmochanes hemileucus* | 21 | 2,358 | 5,910 | | 6,918 | 722 |
| *Myrmotherula klagesi* | 19 | 2,353 | 19,424 | | 14,514 | 1,458 |
| *Myrmotherula assimilis* | 38 | 2,355 | 27,670 | | 23,569 | 1,555 |
| *Thamnophilus cryptoleucus* | 22 | 2,347 | 28,632 | | 21,453 | 1,059 |
| *Myrmoborus lugubris* | 80 | 2,358 | 25,944 | | 19,622 | 1,335 |

1 - From a probe set of 2,321 UCEs and 96 exons.

**Table S3.** Contribution of total variation from each genetic cluster (*K* = 1–6) to the best-fit spatial model for *Conirostrum bicolor*. A threshold of 0.02 was used to determine significant contributions.

| K1 | K2 | K3 | K4 | K5 | K6 |
| --- | --- | --- | --- | --- | --- |
| 1 | 0.720565 | 0.6629 | 0.713265 | 0.700013 | 0.664323 |
| 0 | 0.279435 | 0.333759 | 0.274674 | 0.282011 | 0.31606 |
| 0 | 0 | 0.00334 | 0.010453 | 0.010091 | 0.012497 |
| 0 | 0 | 0 | 0.001607 | 0.005571 | 0.004985 |
| 0 | 0 | 0 | 0 | 0.002314 | 0.001375 |
| 0 | 0 | 0 | 0 | 0 | 0.000759 |

**Table S4.** Contribution of total variation from each genetic cluster (*K* = 1–6) to the best-fit spatial model for *Dendroplex kienerii*. A threshold of 0.02 was used to determine significant contributions.

| K1 | K2 | K3 | K4 | K5 | K6 |
| --- | --- | --- | --- | --- | --- |
| 1 | 0.791937 | 0.802986 | 0.726966 | 0.791073 | 0.57455 |
| 0 | 0.208063 | 0.184502 | 0.249032 | 0.087608 | 0.189803 |
| 0 | 0 | 0.012512 | 0.023074 | 0.087342 | 0.147218 |
| 0 | 0 | 0 | 0.000929 | 0.018568 | 0.077683 |
| 0 | 0 | 0 | 0 | 0.015408 | 0.007132 |
| 0 | 0 | 0 | 0 | 0 | 0.003614 |

**Table S5**. Contribution of total variation from each genetic cluster (*K* = 1–6) to the best-fit non-spatial model for *Mazaria propinqua*. A threshold of 0.02 was used to determine significant contributions.

| K1 | K2 | K3 | K4 | K5 | K6 |
| --- | --- | --- | --- | --- | --- |
| 1 | 0.701268 | 0.700249 | 0.54106 | 0.3912 | 0.251336 |
| 0 | 0.298732 | 0.285355 | 0.362425 | 0.383352 | 0.381998 |
| 0 | 0 | 0.014396 | 0.05989 | 0.121156 | 0.247031 |
| 0 | 0 | 0 | 0.036625 | 0.103209 | 0.098537 |
| 0 | 0 | 0 | 0 | 0.001082 | 0.018578 |
| 0 | 0 | 0 | 0 | 0 | 0.00252 |

**Table S6.** Contribution of total variation from each genetic cluster (K = 1–6) to the best-fit spatial model for *Cranioleuca vulpecula*. A threshold of 0.02 was used to determine significant contributions.

| K1 | K2 | K3 | K4 | K5 | K6 |
| --- | --- | --- | --- | --- | --- |
| 1 | 0.949214 | 0.933531 | 0.592927 | 0.752393 | 0.478746 |
| 0 | 0.050786 | 0.065722 | 0.275043 | 0.10381 | 0.200032 |
| 0 | 0 | 0.000748 | 0.117067 | 0.088896 | 0.17878 |
| 0 | 0 | 0 | 0.014963 | 0.052657 | 0.138348 |
| 0 | 0 | 0 | 0 | 0.002244 | 0.003271 |
| 0 | 0 | 0 | 0 | 0 | 0.000823 |

**Table S7.** Contribution of total variation from each genetic cluster (K = 1–6) to the best-fit spatial model for *Furnarius minor*. A threshold of 0.02 was used to determine significant contributions.

| K1 | K2 | K3 | K4 | K5 | K6 |
| --- | --- | --- | --- | --- | --- |
| 1 | 0.989973 | 0.982845 | 0.948181 | 0.956942 | 0.973881 |
| 0 | 0.010027 | 0.011689 | 0.032764 | 0.022852 | 0.007957 |
| 0 | 0 | 0.005466 | 0.018585 | 0.013763 | 0.006714 |
| 0 | 0 | 0 | 0.000469 | 0.005461 | 0.00549 |
| 0 | 0 | 0 | 0 | 0.000983 | 0.004031 |
| 0 | 0 | 0 | 0 | 0 | 0.001926 |

**Table S8.** Contribution of total variation from each genetic cluster (K = 1–6) to the best-fit spatial model for *Serpophaga hypoleuca*. A threshold of 0.02 was used to determine significant contributions.

| K1 | K2 | K3 | K4 | K5 | K6 |
| --- | --- | --- | --- | --- | --- |
| 1 | 0.951532 | 0.974761 | 0.970605 | 0.952696 | 0.957799 |
| 0 | 0.038468 | 0.024008 | 0.025183 | 0.023246 | 0.025878 |
| 0 | 0 | 0.001231 | 0.003688 | 0.020335 | 0.004372 |
| 0 | 0 | 0 | 0.000523 | 0.001887 | 0.000964 |
| 0 | 0 | 0 | 0 | 0.001837 | 0.00073 |
| 0 | 0 | 0 | 0 | 0 | 0.000258 |

**Table S9.** Contribution of total variation from each genetic cluster (K = 1–6) to the best-fit spatial model for *Stigmatura napensis*. A threshold of 0.02 was used to determine significant contributions.

| K1 | K2 | K3 | K4 | K5 | K6 |
| --- | --- | --- | --- | --- | --- |
| 1 | 0.718225 | 0.44295 | 0.537323 | 0.390576 | 0.39706 |
| 0 | 0.281775 | 0.333308 | 0.345235 | 0.379102 | 0.273861 |
| 0 | 0 | 0.223742 | 0.113586 | 0.154728 | 0.185489 |
| 0 | 0 | 0 | 0.003855 | 0.074924 | 0.101902 |
| 0 | 0 | 0 | 0 | 0.00067 | 0.039636 |
| 0 | 0 | 0 | 0 | 0 | 0.002052 |

**Table S10.** Contribution of total variation from each genetic cluster (K = 1–6) to the best-fit spatial model for *Myrmochanes hemileucus*. A threshold of 0.02 was used to determine significant contributions.

| K1 | K2 | K3 | K4 | K5 | K6 |
| --- | --- | --- | --- | --- | --- |
| 1 | 0.503788 | 0.501642 | 0.494904 | 0.641486 | 0.539253 |
| 0 | 0.496212 | 0.403635 | 0.447757 | 0.324631 | 0.448794 |
| 0 | 0 | 0.094723 | 0.03384 | 0.022797 | 0.005862 |
| 0 | 0 | 0 | 0.023499 | 0.008737 | 0.002708 |
| 0 | 0 | 0 | 0 | 0.002349 | 0.002317 |
| 0 | 0 | 0 | 0 | 0 | 0.001065 |

**Table S11.** Contribution of total variation from each genetic cluster (K = 1–6) to the best-fit spatial model for *Myrmotherula klagesi*. A threshold of 0.02 was used to determine significant contributions.

| K1 | K2 | K3 | K4 | K5 | K6 |
| --- | --- | --- | --- | --- | --- |
| 1 | 0.522816 | 0.579243 | 0.514522 | 0.498874 | 0.515386 |
| 0 | 0.477184 | 0.220051 | 0.282525 | 0.295399 | 0.272208 |
| 0 | 0 | 0.200706 | 0.200486 | 0.200538 | 0.200269 |
| 0 | 0 | 0 | 0.002467 | 0.002682 | 0.000654 |
| 0 | 0 | 0 | 0 | 0.002507 | 0.002862 |
| 0 | 0 | 0 | 0 | 0 | 0.008621 |

**Table S12.** Contribution of total variation from each genetic cluster (K = 1–6) to the best-fit spatial model for *Myrmotherula assimilis*. A threshold of 0.02 was used to determine significant contributions.

| K1 | K2 | K3 | K4 | K5 | K6 |
| --- | --- | --- | --- | --- | --- |
| 1 | 0.300676 | 0.000231 | 0.083514 | 0.002636 | 0.000756 |
| 0 | 0.699324 | 0.696768 | 0.701513 | 0.776121 | 0.000287 |
| 0 | 0 | 0.303001 | 0.027781 | 0.000144 | 0.200645 |
| 0 | 0 | 0 | 0.187191 | 0.217611 | 0.000371 |
| 0 | 0 | 0 | 0 | 0.003499 | 0.000308 |
| 0 | 0 | 0 | 0 | 0 | 0.797634 |

**Table S13.** Contribution of total variation from each genetic cluster (K = 1–6) to the best-fit spatial model for *Thamnophilus cryptoleucus*. A threshold of 0.02 was used to determine significant contributions.

| K1 | K2 | K3 | K4 | K5 | K6 |
| --- | --- | --- | --- | --- | --- |
| 1 | 0.888177 | 0.944532 | 0.940586 | 0.935745 | 0.952381 |
| 0 | 0.111823 | 0.055287 | 0.055177 | 0.041837 | 0.037292 |
| 0 | 0 | 0.010181 | 0.002173 | 0.009528 | 0.003354 |
| 0 | 0 | 0 | 0.002064 | 0.006762 | 0.002738 |
| 0 | 0 | 0 | 0 | 0.006133 | 0.002405 |
| 0 | 0 | 0 | 0 | 0 | 0.001911 |

**Table S14.** Contribution of total variation from each genetic cluster (K = 1–6) to the best-fit spatial model for *Myrmoborus lugubris*. A threshold of 0.02 was used to determine significant contributions.

| K1 | K2 | K3 | K4 | K5 | K6 |
| --- | --- | --- | --- | --- | --- |
| 1 | 0.726634 | 0.702905 | 0.714519 | 0.705926 | 0.693205 |
| 0 | 0.273366 | 0.29527 | 0.281506 | 0.289818 | 0.293237 |
| 0 | 0 | 0.001825 | 0.002606 | 0.001746 | 0.007109 |
| 0 | 0 | 0 | 0.001369 | 0.001433 | 0.002627 |
| 0 | 0 | 0 | 0 | 0.001084 | 0.002172 |
| 0 | 0 | 0 | 0 | 0 | 0.001722 |

**Table S15.** Weir and Cockerham’s *F*_ST_ values between populations within each species of Amazonian riverine island birds. *Furnarius minor* was excluded due to a single inferred population (*K* = 1).

| Species | Sub-basin population comparison | Weir and Cockerham’s *F*_ST_ |
| --- | --- | --- |
| *Conirostrum bicolor* | Amazonas – Branco | 0.106 |
| *Dendroplex kienerii* | Amazonas – Solimões | 0.063 |
| *Mazaria propinqua* | Amazonas – Branco | 0.03 |
| *Cranioleuca vulpecula* | Solimões – Madeira | 0.078 |
| *Serpophaga hypoleuca* | Amazonas – Branco | 0.062 |
| *Stigmatura napensis* | Amazonas – Branco | 0.041 |
| *Myrmochanes hemileucus* | Solimões – Ucayali | 0.086 |
| *Myrmotherula klagesi* | Solimões – Branco | 0.106 |
|  | Amazonas – Branco | 0.093 |
|  | Amazonas – Solimões | 0.053 |
| *Myrmotherula assimilis* | Amazonas – Madeira | 0.166 |
| *Thamnophilus cryptoleucus* | Solimões – Madeira | 0.116 |
| *Myrmoborus lugubris* | Amazonas – Solimões | 0.258 |

**Table S16.** Genetic diversity metrics for populations grouped by Amazonian sub-basins, including nucleotide diversity (π), observed heterozygosity (Ho), and Tajima’s D.

| Species | Population / sub-basin | N | π | Ho | Tajima`s *D* |
| --- | --- | --- | --- | --- | --- |
| *Conirostrum bicolor* | Amazonas | 9 | 0.00173 | 0.165 | -0.428 |
|  | Branco | 6 | 0.00154 | 0.1581 | -0.579 |
| *Dendrople kienerii* | Amazonas | 15 | 0.00262 | 0.1463 | 0.319 |
|  | Solimões | 4 | 0.00218 | 0.1538 | 0.151 |
| *Mazaria propinqua* | Amazonas | 16 | 0.00186 | 0.1755 | -0.365 |
|  | Branco | 7 | 0.00145 | 0.1568 | -0.217 |
| *Cranioleuca vulpecula* | Amazonas | 22 | 0.00238 | 0.1785 | 0.098 |
|  | Madeira | 4 | 0.00172 | 0.1829 | -0.168 |
| *Furnarius minor* | Amazonas | 10 | 0.00173 | 0.1683 | 0.284 |
| *Serpophaga hypoleuca* | Amazonas | 14 | 0.00258 | 0.1755 | 0.064 |
|  | Branco | 2 | 0.00174 | 0.1758 | 0.364 |
| *Stigmatura napensis* | Amazonas | 22 | 0.00242 | 0.1682 | 0.157 |
|  | Branco | 7 | 0.00156 | 0.1618 | 0.407 |
| *Myrmotherula hemileucus* | Solimões | 18 | 0.00266 | 0.2524 | 0.205 |
|  | Ucayali | 3 | 0.00198 | 0.1971 | 0.336 |
| *Myrmotherula klagesi* | Amazonas | 7 | 0.00167 | 0.1538 | -0.431 |
|  | Solimões | 4 | 0.00149 | 0.1401 | -0.217 |
|  | Branco | 9 | 0.00162 | 0.1463 | -0.378 |
| *Myrmotherula assimilis* | Solimões | 13 | 0.00163 | 0.1425 | -0.243 |
|  | Madeira | 25 | 0.00138 | 0.1389 | -0.308 |
| *Thamnophilus cryptoleucus* | Solimões | 19 | 0.00156 | 0.1461 | -0.607 |
|  | Madeira | 3 | 0.00143 | 0.1848 | -0.088 |
| *Myrmoborus lugubris* | Amazonas | 31 | 0.00166 | 0.0716 | -0.612 |
|  | Solimões | 49 | 0.00181 | 0.0944 | -0.564 |

**Table S17.** Estimated demographic parameters under the best-fit isolation with secondary contact model for *Conirostrum bicolor*, including demographic units, prior distributions, and estimates based on a mutation rate of 2.5 × 10⁻⁹ with 95% confidence intervals.

| Parameters | Demographic unit | Prior distribution | Estimate based on *μ* = 2.5 x 10^-9^ (CI 95%) |
| --- | --- | --- | --- |
| Ne_A_ | Current population size – Amazonas | 100 – 50,000 | 51,133 (17,354 – 80,823) |
| Ne_B_ | Current population size – Branco | 100 – 50,000 | 3,461 (1,290 – 17,624) |
| Ne_AB_ | Ancestral population size | 1,000 – 500,000 | 12, 062 (3,822 – 89,016) |
| Tdiv_AB_ | Divergence time | 10,000 – 500,000 | 21,786 (10,711 – 49,200) |
| Tsc_AB_ | Time of secondary contact | 1,000 – 50,000 | 7,440 (5,794 – 18,001) |
| m_AB_ | Migration from Amazonas to Branco | 0.001 – 10 | 0.161 (0.004 – 1.744) |
| m_BA_ | Migration from Branco to Amazonas | 0.001 - 10 | 1.895 (0.098 – 3.714) |

**Table S18.** Estimated demographic parameters under the best-fit isolation with secondary contact model for *Dendroplex kienerii*, including demographic units, prior distributions, and estimates based on a mutation rate of 2.5 × 10⁻⁹ with 95% confidence intervals.

| Parameters | Demographic unit | Prior distribution | Estimate based on *μ* = 2.5 x 10^-9^ (CI 95%) |
| --- | --- | --- | --- |
| Ne_A_ | Current population size – Amazonas | 100 – 50,000 | 1,168 (3,835 – 85,102) |
| Ne_B_ | Current population size – Solimões | 100 – 50,000 | 5,522 (2,973 – 43,824) |
| Ne_AB_ | Ancestral population size | 1,000 – 500,000 | 277,514 (102,688 – 291,720) |
| Tdiv_AB_ | Divergence time | 10,000 – 500,000 | 4,158 (14,290 – 40,762) |
| Tsc_AB_ | Time of secondary contact | 1,000 – 50,000 | 4,005 (2,873 – 21,832) |
| m_AB_ | Migration from Amazonas to Solimões | 0.001 – 10 | 0.589 (0.030 – 1.517) |
| m_BA_ | Migration from Solimões to Amazonas | 0.001 - 10 | 3.428 (0.753 – 6.109) |

**Table S19.** Estimated demographic parameters under the best-fit isolation with secondary contact model for *Mazaria propinqua*, including demographic units, prior distributions, and estimates based on a mutation rate of 2.5 × 10⁻⁹ with 95% confidence intervals.

| Parameters | Demographic unit | Prior distribution | Estimate based on *μ* = 2.5 x 10^-9^ (CI 95%) |
| --- | --- | --- | --- |
| Ne_A_ | Current population size – Amazonas | 100 – 50,000 | 84,172 (44,933 – 121,706) |
| Ne_B_ | Current population size – Branco | 100 – 50,000 | 2,151 (813 – 36,995) |
| Ne_AB_ | Ancestral population size | 1,000 – 500,000 | 22,723 (7,296 – 72,511) |
| Tdiv_AB_ | Divergence time | 10,000 – 500,000 | 4,042 (708 – 40,351) |
| Tsc_AB_ | Time of secondary contact | 1,000 – 50,000 | 809 (2,291 – 18,006) |
| m_AB_ | Migration from Amazonas to Branco | 0.001 – 10 | 0.029 (0,004 – 0.394) |
| m_BA_ | Migration from Branco to Amazonas | 0.001 - 10 | 6.036 (2.571 – 7.404) |

**Table S20.** Estimated demographic parameters under the best-fit isolation with secondary contact model for *Cranioleuca vulpecula*, including demographic units, prior distributions, and estimates based on a mutation rate of 2.5 × 10⁻⁹ with 95% confidence intervals.

| Parameters | Demographic unit | Prior distribution | Estimate based on *μ* = 2.5 x 10^-9^ (CI 95%) |
| --- | --- | --- | --- |
| Ne_A_ | Current population size – Solimões | 100 – 50,000 | 10,027 (7,291 – 64,292) |
| Ne_B_ | Current population size – Madeira | 100 – 50,000 | 1,720 (4,054 – 36,266) |
| Ne_AB_ | Ancestral population size | 1,000 – 500,000 | 99,088 (32,982 – 216,067) |
| Tdiv_AB_ | Divergence time | 10,000 – 500,000 | 2,206 (1,923 – 34,032) |
| Tsc_AB_ | Time of secondary contact | 1,000 – 50,000 | 1,806 (928 – 28,054) |
| m_AB_ | Migration from Solimões to Madeira | 0.001 – 10 | 1.904 (0.731 – 3.088) |
| m_BA_ | Migration from Madeira to Solimões | 0.001 - 10 | 1.240 (0.394 – 2.308) |

**Table S21.** Estimated current effective population size for *Furnarius minor*, including prior distributions, and estimates based on a mutation rate of 2.5 × 10⁻⁹ with 95% confidence intervals.

| Parameters | Demographic unit | Prior distribution | Estimate based on *μ* = 2.5 x 10^-9^ (CI 95%) |
| --- | --- | --- | --- |
| Ne | Current population size - Amazonas | 100 – 50,000 | 33,391 (7,978 – 102,855) |

**Table S22.** Estimated demographic parameters under the best-fit isolation with secondary contact model for *Serpophaga hypoleuca*, including demographic units, prior distributions, and estimates based on a mutation rate of 2.5 × 10⁻⁹ with 95% confidence intervals.

| Parameters | Demographic unit | Prior distribution | Estimate based on *μ* = 2.5 x 10^-9^ (CI 95%) |
| --- | --- | --- | --- |
| Ne_A_ | Current population size – Amazonas | 100 – 50,000 | 32,716 (28,640 – 37, 901) |
| Ne_B_ | Current population size – Branco | 100 – 50,000 | 2,432 (1,202 – 6,188) |
| Ne_AB_ | Ancestral population size | 1,000 – 500,000 | 8,155 (6,631 – 17,394) |
| Tdiv_AB_ | Divergence time | 10,000 – 500,000 | 2,566 (2,307 – 34,724) |
| Tsc_AB_ | Time of secondary contact | 1,000 – 50,000 | 1,745 (1,006 – 18,176) |
| m_AB_ | Migration from Amazonas to Branco | 0.001 – 10 | 0.949 (0.266 – 2.398) |
| m_BA_ | Migration from Branco to Amazonas | 0.001 - 10 | 3.469 (1.734 – 5.006) |

**Table S23.** Estimated demographic parameters under the best-fit isolation with secondary contact model for *Stigmatura napensis*, including demographic units, prior distributions, and estimates based on a mutation rate of 2.5 × 10⁻⁹ with 95% confidence intervals.

| Parameters | Demographic unit | Prior distribution | Estimate based on *μ* = 2.5 x 10^-9^ (CI 95%) |
| --- | --- | --- | --- |
| Ne_A_ | Current population size – Amazonas | 100 – 50,000 | 7,570 (3,855 – 39,712) |
| Ne_B_ | Current population size – Branco | 100 – 50,000 | 179 (44 – 5,046) |
| Ne_AB_ | Ancestral population size | 1,000 – 500,000 | 1,821 (617 – 39,007) |
| Tdiv_AB_ | Divergence time | 10,000 – 500,000 | 7,284 (1,271 – 59, 443) |
| Tsc_AB_ | Time of secondary contact | 1,000 – 50,000 | 1,828 (729 – 40,513) |
| m_AB_ | Migration from Amazonas to Branco | 0.001 – 10 | 0.268 (0.073 – 1.404) |
| m_BA_ | Migration from Branco to Amazonas | 0.001 - 10 | 5.850 (3.362 – 6.777) |

**Table S24.** Estimated demographic parameters under the best-fit isolation with secondary contact model for *Myrmochanes hemileucus*, including demographic units, prior distributions, and estimates based on a mutation rate of 2.5 × 10⁻⁹ with 95% confidence intervals.

| Parameters | Demographic unit | Prior distribution | Estimate based on *μ* = 2.5 x 10^-9^ (CI 95%) |
| --- | --- | --- | --- |
| Ne_A_ | Current population size – Solimões | 100 – 50,000 | 38,882 (17,927 – 106,383) |
| Ne_B_ | Current population size – Ucayali | 100 – 50,000 | 31,648 (28,005 – 66,817) |
| Ne_AB_ | Ancestral population size | 1,000 – 500,000 | 217,797 (167,267 – 370,602) |
| Tdiv_AB_ | Divergence time | 10,000 – 500,000 | 50,174 (23,409 – 83,545) |
| Tsc_AB_ | Time of secondary contact | 1,000 – 50,000 | 16,922 (13,005 – 41,941) |
| m_AB_ | Migration from Solimões to Ucayali | 0.001 – 10 | 2.041 (0.881 – 3.932) |
| m_BA_ | Migration from Ucayali to Solimões | 0.001 - 10 | 0.027 (0.006 – 1.062) |

**Table S25.** Estimated demographic parameters under the best-fit isolation with secondary contact model for *Myrmotherula klagesi*, including demographic units, prior distributions, and estimates based on a mutation rate of 2.5 × 10⁻⁹ with 95% confidence intervals.

| Parameters | Demographic unit | Prior distribution | Estimate based on *μ* = 2.5 x 10^-9^ (CI 95%) |
| --- | --- | --- | --- |
| Ne_A_ | Current population size – Amazonas | 100 – 50,000 | 296,587 (202,754 – 386,220) |
| Ne_B_ | Current population size – Solimões | 100 – 50,000 | 146,753 (59,129 – 187,300) |
| Ne_C_ | Current population size – Branco | 100 – 50,000 | 166,296 (18,722 – 237,371) |
| Ne_AB_ | Ancestral population size Amazonas-Branco | 1,000 – 500,000 | 163,446 (102,805 – 209,267) |
| Ne_ABC_ | Ancestral population size Amanonas-Solimões-Branco | 1,000 – 500,000 | 223,240 (202,011 – 332,844) |
| Tdiv_AB_ | Divergence time between Amazonas-Solimões | 10,000 – 500,000 | 174,808 (148,218 – 220,669) |
| Tdiv_AC_ | Divergence time between Amazonas-Branco | 10,000 – 500,000 | 257,289 (215,932 – 287,050) |
| Tsc_AB_ | Time of secondary contact Amazonas-Solimões | 1,000 – 50,000 | 18,478 (6,297 – 44,843) |
| Tsc_AC_ | Time of secondary contact Amazonas-Branco | 1,000 – 50,000 | 56,220 (18,091 – 103,653) |
| m_AB_ | Migration from Amazonas to Solimões | 0.001 – 10 | 2.130 (0.219 – 3.142) |
| m_BA_ | Migration from Solimões to Amazonas | 0.001 - 10 | 3.288 (1.004 – 3.142) |
| m_AC_ | Migration from Amazonas to Branco | 0.001 – 10 | 1.036 (0.512 – 2.985) |
| m_CA_ | Migration from Branco to Amazonas | 0.001 - 10 | 2.650 (1.613 – 4.046) |

**Table S26.** Estimated demographic parameters under the best-fit isolation without gene flow model for *Myrmotherula assimilis*, including demographic units, prior distributions, and estimates based on a mutation rate of 2.5 × 10⁻⁹ with 95% confidence intervals.

| Parameters | Demographic unit | Prior distribution | Estimate based on *μ* = 2.5 x 10^-9^ (CI 95%) |
| --- | --- | --- | --- |
| Ne_A_ | Current population size – Amazonas | 100 – 50,000 | 20,779 (6,883 – 41,005) |
| Ne_B_ | Current population size – Madeira | 100 – 50,000 | 10,176 (2,931 – 37,273) |
| Ne_AB_ | Ancestral population size | 1,000 – 500,000 | 306,983 (200,804 – 317,436) |
| Tdiv_AB_ | Divergence time | 10,000 – 500,000 | 22,594 (10,872 – 44,332) |

**Table S27.** Estimated demographic parameters under the best-fit isolation with secondary contact model for *Thamnophilus cryptoleucus*, including demographic units, prior distributions, and estimates based on a mutation rate of 2.5 × 10⁻⁹ with 95% confidence intervals.

| Parameters | Demographic unit | Prior distribution | Estimate based on *μ* = 2.5 x 10^-9^ (CI 95%) |
| --- | --- | --- | --- |
| Ne_A_ | Current population size – Solimões | 100 – 50,000 | 313,920 (287,832 – 370,302) |
| Ne_B_ | Current population size – Madeira | 100 – 50,000 | 312,851 (269,301 – 333,276) |
| Ne_AB_ | Ancestral population size | 1,000 – 500,000 | 297,451(273,734 – 439,079) |
| Tdiv_AB_ | Divergence time | 10,000 – 500,000 | 120,234 (87,832 – 158,934) |
| Tsc_AB_ | Time of secondary contact | 1,000 – 50,000 | 64,988 (50,611 – 89,437) |
| m_AB_ | Migration from Solimões to Madeira | 0.001 – 10 | 1.596 (0.085 – 2.003) |
| m_BA_ | Migration from Madeira to Solimões | 0.001 - 10 | 0.920 (0.052 – 2.474) |

**Table S28.** Estimated demographic parameters under the best-fit isolation with secondary contact model for *Myrmoborus lugubris*, including demographic units, prior distributions, and estimates based on a mutation rate of 2.5 × 10⁻⁹ with 95% confidence intervals.

| Parameters | Demographic unit | Prior distribution | Estimate based on *μ* = 2.5 x 10^-9^ (CI 95%) |
| --- | --- | --- | --- |
| Ne_A_ | Current population size – Amazonas | 100 – 50,000 | 53,051 (40,465 – 85,113) |
| Ne_B_ | Current population size – Solimões | 100 – 50,000 | 19,058 (18,332 – 25,298) |
| Ne_AB_ | Ancestral population size | 1,000 – 500,000 | 7,942 (5,356 – 33,824) |
| Tdiv_AB_ | Divergence time | 10,000 – 500,000 | 58,406 (42,832 – 94,060) |
| Tsc_AB_ | Time of secondary contact | 1,000 – 50,000 | 23,927 (7,093 – 62,144) |
| m_AB_ | Migration from Amazonas to Solimões | 0.001 – 10 | 0.296 (0.006 – 1.825) |
| m_BA_ | Migration from Solimões to Amazonas | 0.001 - 10 | 0.221 (0.090 – 1.467) |

**Table S29.** Results of the Bayesian phylogenetic multilevel model with *F*_ST_ as the response variable.

| **Parameters** | **Estimate** | **SE** | **L-95%** | **U-95%** | **pMCMC** | **Rhat** | **Bulk ESS** | **Tail ESS** | **R^2^** |
| --- | --- | --- | --- | --- | --- | --- | --- | --- | --- |
| Intercept | 0.08 | 0.19 | -0.28 | 0.48 | 0.689 | 1.00 | 11582 | 12463 | 0.20 |
| Vegetation: Shrub – Forest | **-0.77** | **0.25** | **-1.28** | **-0.31** | **< 0.001** | 1.00 | 14433 | 15000 |  |

Legend: Est. Error is the estimated error, L- and U-95% are lower and upper bounds of 95% credible intervals, respectively, pMPE is the p-value obtained from the Bayesian maximum probability of effect, $\hat{R}$ is the Gelman-Rubin diagnostic, bulk and tail ESS are effective sample sizes, and R^2^ is the coefficient of determination. Values in bold indicate effects that are credibly different from zero.

**Table S30.** Results of the Bayesian phylogenetic multilevel model with migration rate as the response variable.

| **Parameters** | **Estimate** | **SE** | **L-95%** | **U-95%** | **pMCMC** | **Rhat** | **Bulk ESS** | **Tail ESS** | **R^2^** |
| --- | --- | --- | --- | --- | --- | --- | --- | --- | --- |
| Intercept | -0.39 | 0.3 | -0.99 | 0.16 | 0.186 | 1.00 | 9763 | 11597 | 0.33 |
| Vegetation: Shrub – Forest | **1.02** | **0.27** | **0.49** | **1.54** | **< 0.001** | 1.00 | 17199 | 15349 |  |

Legend: Est. Error is the estimated error, L- and U-95% are lower and upper bounds of 95% credible intervals, respectively, pMPE is the p-value obtained from the Bayesian maximum probability of effect, $\hat{R}$ is the Gelman-Rubin diagnostic, bulk and tail ESS are effective sample sizes, and R^2^ is the coefficient of determination. Values in bold indicate effects that are credibly different from zero.

**Table S31.** Results of the Bayesian phylogenetic multilevel model with divergence time as the response variable.

| **Parameters** | **Estimate** | **SE** | **L-95%** | **U-95%** | **pMCMC** | **Rhat** | **Bulk ESS** | **Tail ESS** | **R^2^** |
| --- | --- | --- | --- | --- | --- | --- | --- | --- | --- |
| Intercept | -0.10 | 0.15 | -0.38 | 0.20 | 0.48 | 1.00 | 16363 | 14139 | 0.11 |
| Vegetation: Shrub - Forest | **-0.45** | **0.23** | **-0.95** | **-0.05** | **0.03** | 1.00 | 17245 | 14135 |  |

Legend: Est. Error is the estimated error, L- and U-95% are lower and upper bounds of 95% credible intervals, respectively, pMPE is the p-value obtained from the Bayesian maximum probability of effect, $\hat{R}$ is the Gelman-Rubin diagnostic, bulk and tail ESS are effective sample sizes, and R^2^ is the coefficient of determination. Values in bold indicate effects that are credibly different from zero.

**Table S32.** Results of the Bayesian phylogenetic multilevel model with time since secondary contact (Tsc) as the response variable.

| **Parameters** | **Estimate** | **SE** | **L-95%** | **U-95%** | **pMCMC** | **Rhat** | **Bulk ESS** | **Tail ESS** | **R^2^** |
| --- | --- | --- | --- | --- | --- | --- | --- | --- | --- |
| Intercept | 0.02 | 0.17 | -0.29 | 0.38 | 0.92 | 1.00 | 10518 | 11063 | 0.21 |
| Vegetation: Shrub - Forest | **-0.57** | **0.23** | **-1.06** | **-0.16** | **0.01** | 1.00 | 12512 | 11927 |  |

Legend: Est. Error is the estimated error, L- and U-95% are lower and upper bounds of 95% credible intervals, respectively, pMPE is the p-value obtained from the Bayesian maximum probability of effect, $\hat{R}$ is the Gelman-Rubin diagnostic, bulk and tail ESS are effective sample sizes, and R^2^ is the coefficient of determination. Values in bold indicate effects that are credibly different from zero.

**Table S33.** Results of the Bayesian phylogenetic multilevel model with effective population size (Ne) as the response variable.

| **Parameters** | **Estimate** | **SE** | **L-95%** | **U-95%** | **pMCMC** | **Rhat** | **Bulk ESS** | **Tail ESS** | **R^2^** |
| --- | --- | --- | --- | --- | --- | --- | --- | --- | --- |
| Intercept | 0.27 | 0.21 | -0.13 | 0.69 | 0.20 | 1.00 | 5999 | 9628 | 0.72 |
| Vegetation: Shrub - Forest | **-0.56** | **0.25** | **-1.06** | **-0.08** | **0.02** | 1.00 | 6596 | 10457 |  |
| Basin: Branco - Amazonas | **-0.34** | **0.16** | **-0.65** | **-0.01** | **0.04** | 1.00 | 13441 | 10802 |  |
| Basin: Amazonas - Madeira | -0.11 | 0.2 | -0.5 | 0.28 | 0.55 | 1.00 | 15296 | 13475 |  |
| Basin: Solimões - Amazonas | -0.19 | 0.16 | -0.49 | 0.13 | 0.25 | 1.00 | 10937 | 12247 |  |

Legend: Est. Error is the estimated error, L- and U-95% are lower and upper bounds of 95% credible intervals, respectively, pMPE is the p-value obtained from the Bayesian maximum probability of effect, $\hat{R}$ is the Gelman-Rubin diagnostic, bulk and tail ESS are effective sample sizes, and R^2^ is the coefficient of determination. Values in bold indicate effects that are credibly different from zero.

**Table S34.** Contrasts between hydrological basins in the model with effective population size (Ne) as the response variable.

| **Contrast** | **Estimate** | **L-95%** | **U-95%** | **pMCMC** |
| --- | --- | --- | --- | --- |
| Amazonas - Branco | **0.34** | **0.02** | **0.65** | **0.04** |
| Amazonas - Madeira | 0.11 | -0.28 | 0.49 | 0.55 |
| Amazonas - Solimões | 0.19 | -0.13 | 0.49 | 0.25 |
| Branco - Madeira | -0.23 | -0.67 | 0.25 | 0.32 |
| Branco – Solimões | -0.15 | -0.54 | 0.22 | 0.43 |
| Madeira - Solimões | 0.07 | -0.31 | 0.46 | 0.69 |

Legend: L- and U-95% are lower and upper bounds of 95% credible intervals.

**Table S35.** Results of the Bayesian phylogenetic multilevel model with nucleotide diversity (π) as the response variable.

| **Parameters** | **Estimate** | **Est. Error** | **L-95%** | **U-95%** | **pMCMC** | **Rhat** | **Bulk ESS** | **Tail ESS** | **R^2^** |
| --- | --- | --- | --- | --- | --- | --- | --- | --- | --- |
| Intercept | -0.03 | 0.19 | -0.42 | 0.32 | 0.88 | 1.00 | 14069 | 14050 | 0.36 |
| Vegetation: Shrub - Forest | **0.54** | **0.22** | **0.12** | **0.98** | **0.01** | 1.00 | 14212 | 13592 |  |
| Basin: Branco - Amazonas | **-0.92** | **0.20** | **-1.32** | **-0.52** | **< 0.001** | 1.00 | 17506 | 14525 |  |
| Basin: Amazonas - Madeira | **-0.78** | **0.26** | **-1.32** | **-0.31** | **0.002** | 1.00 | 15540 | 14351 |  |
| Basin: Solimões - Amazonas | -0.17 | 0.21 | -0.55 | 0.28 | 0.41 | 1.00 | 15370 | 13752 |  |

Legend: Est. Error is the estimated error, L- and U-95% are lower and upper bounds of 95% credible intervals, respectively, pMPE is the p-value obtained from the Bayesian maximum probability of effect, $\hat{R}$ is the Gelman-Rubin diagnostic, bulk and tail ESS are effective sample sizes, and R^2^ is the coefficient of determination. Values in bold indicate effects that are credibly different from zero.

**Table S36.** Contrasts between hydrological basins in the model with nucleotide diversity (π) as the response variable.

| **Contrast** | **Estimate** | **L-95%** | **U-95%** | **pMPE** |
| --- | --- | --- | --- | --- |
| Amazonas - Branco | **0.92** | **0.52** | **1.31** | **< 0.001** |
| Amazonas - Madeira | **0.77** | **0.28** | **1.29** | **0.002** |
| Amazonas - Solimões | 0.17 | -0.25 | 0.57 | 0.409 |
| Branco - Madeira | -0.15 | -0.70 | 0.46 | 0.621 |
| Branco – Solimões | **-0.75** | **-1.29** | **-0.25** | **0.004** |
| Madeira - Solimões | **-0.59** | **-1.15** | **-0.14** | **0.006** |

Legend: L- and U-95% are lower and upper bounds of 95% credible intervals.

**Table S37.** Results of the Bayesian phylogenetic multilevel model with observed heterozygosity (Ho) as the response variable.

| **Parameters** | **Estimate** | **SE** | **L-95%** | **U-95%** | **pMCMC** | **Rhat** | **Bulk ESS** | **Tail ESS** | **R^2^** |
| --- | --- | --- | --- | --- | --- | --- | --- | --- | --- |
| Intercept | -0.47 | 0.21 | -0.89 | -0.07 | 0.024 | 1.00 | 9493 | 11680 | 0.84 |
| Vegetation: Shrub - Forest | **1.22** | **0.25** | **0.72** | **1.71** | **< 0.001** | 1.00 | 9992 | 13281 |  |
| Basin: Branco - Amazonas | -0.2 | 0.15 | -0.5 | 0.11 | 0.194 | 1.00 | 17231 | 10821 |  |
| Basin: Amazonas - Madeira | 0.33 | 0.25 | -0.14 | 0.85 | 0.163 | 1.00 | 10214 | 10922 |  |
| Basin: Solimões - Amazonas | -0.11 | 0.22 | -0.5 | 0.37 | 0.583 | 1.00 | 7902 | 8547 |  |

Legend: Est. Error is the estimated error, L- and U-95% are lower and upper bounds of 95% credible intervals, respectively, pMPE is the p-value obtained from the Bayesian maximum probability of effect, $\hat{R}$ is the Gelman-Rubin diagnostic, bulk and tail ESS are effective sample sizes, and R^2^ is the coefficient of determination. Values in bold indicate effects that are credibly different from zero.

**Table S38.** Contrasts between hydrological basins in the model with nucleotide diversity (π) as the response variable.

| **Contrast** | **Estimate** | **L-95%** | **U-95%** | **pMPE** |
| --- | --- | --- | --- | --- |
| Amazonas - Branco | 0.20 | -0.10 | 0.50 | 0.19 |
| Amazonas - Madeira | -0.32 | -0.84 | 0.15 | 0.16 |
| Amazonas - Solimões | 0.12 | -0.34 | 0.53 | 0.58 |
| Branco - Madeira | -0.52 | -1.07 | 0.00 | 0.05 |
| Branco – Solimões | -0.08 | -0.57 | 0.38 | 0.73 |
| Madeira - Solimões | 0.44 | -0.01 | 0.86 | 0.06 |

Legend: L- and U-95% are lower and upper bounds of 95% credible intervals.

**Table S39.** Results of the Bayesian phylogenetic multilevel model with Tajima’s D as the response variable.

| **Parameters** | **Estimate** | **SE** | **L-95%** | **U-95%** | **pMCMC** | **Rhat** | **Bulk ESS** | **Tail ESS** | **R^2^** |
| --- | --- | --- | --- | --- | --- | --- | --- | --- | --- |
| Intercept | -0.56 | 0.22 | -0.99 | -0.11 | 0.018 | 1.00 | 15164 | 13373 | 0.54 |
| Vegetation: Shrub - Forest | **1.08** | **0.24** | **0.61** | **1.54** | **< 0.001** | 1.00 | 19245 | 15093 |  |

Legend: Est. Error is the estimated error, L- and U-95% are lower and upper bounds of 95% credible intervals, respectively, pMPE is the p-value obtained from the Bayesian maximum probability of effect, $\hat{R}$ is the Gelman-Rubin diagnostic, bulk and tail ESS are effective sample sizes, and R^2^ is the coefficient of determination. Values in bold indicate effects that are credibly different from zero.

**Table S40.** Results of the Bayesian phylogenetic multilevel model with genetic-geographic similarity (t0) as the response variable.

| **Parameters** | **Estimate** | **SE** | **L-95%** | **U-95%** | **pMCMC** | **Rhat** | **Bulk ESS** | **Tail ESS** | **R^2^** |
| --- | --- | --- | --- | --- | --- | --- | --- | --- | --- |
| Intercept | 0.22 | 0.18 | -0.12 | 0.58 | 0.21 | 1.00 | 15991 | 13977 | 0.29 |
| Vegetation: Shrub – Forest | **-0.87** | **0.2** | **-1.25** | **-0.45** | **< 0.001** | 1.00 | 16877 | 14362 |  |

Legend: Est. Error is the estimated error, L- and U-95% are lower and upper bounds of 95% credible intervals, respectively, pMPE is the p-value obtained from the Bayesian maximum probability of effect, $\hat{R}$ is the Gelman-Rubin diagnostic, bulk and tail ESS are effective sample sizes, and R^2^ is the coefficient of determination. Values in bold indicate effects that are credibly different from zero.

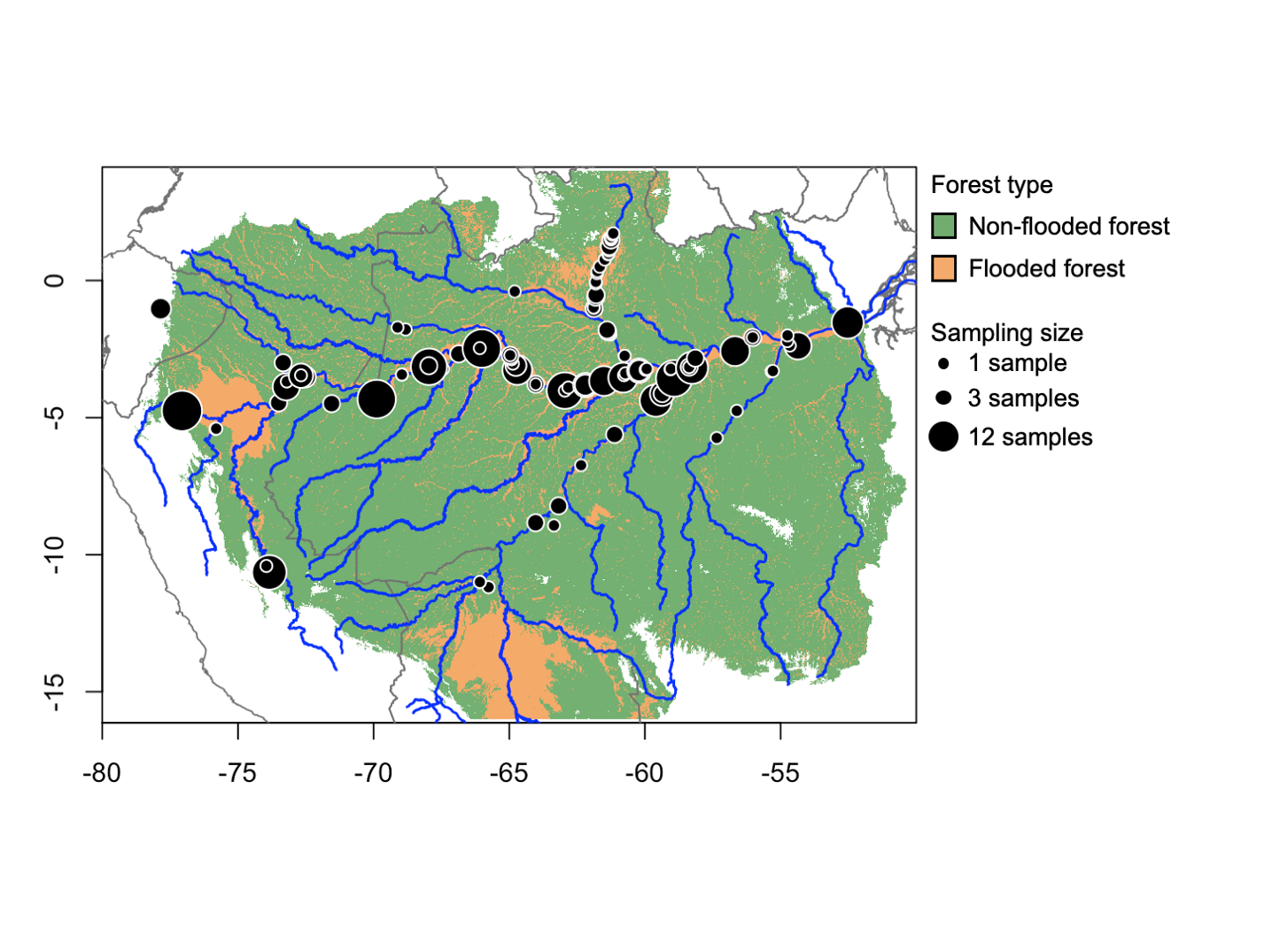

**Figure S1.** Map distribution of samples.

**Figure S2.** Cross-validation results from *conStruct* analyses comparing spatial and non-spatial models and identifying the optimal number of populations (*K*) under the best-fitting model for each of the 12 Amazonian riverine island bird species (panels a–l). See also Table S2.

**Figure S3.** Optimal number of populations (*K*), genetic–geographic similarity (*t₀*) derived from Procrustes analysis, and associated p-values for 12 bird species inhabiting Amazonian riverine islands (panels a–l).
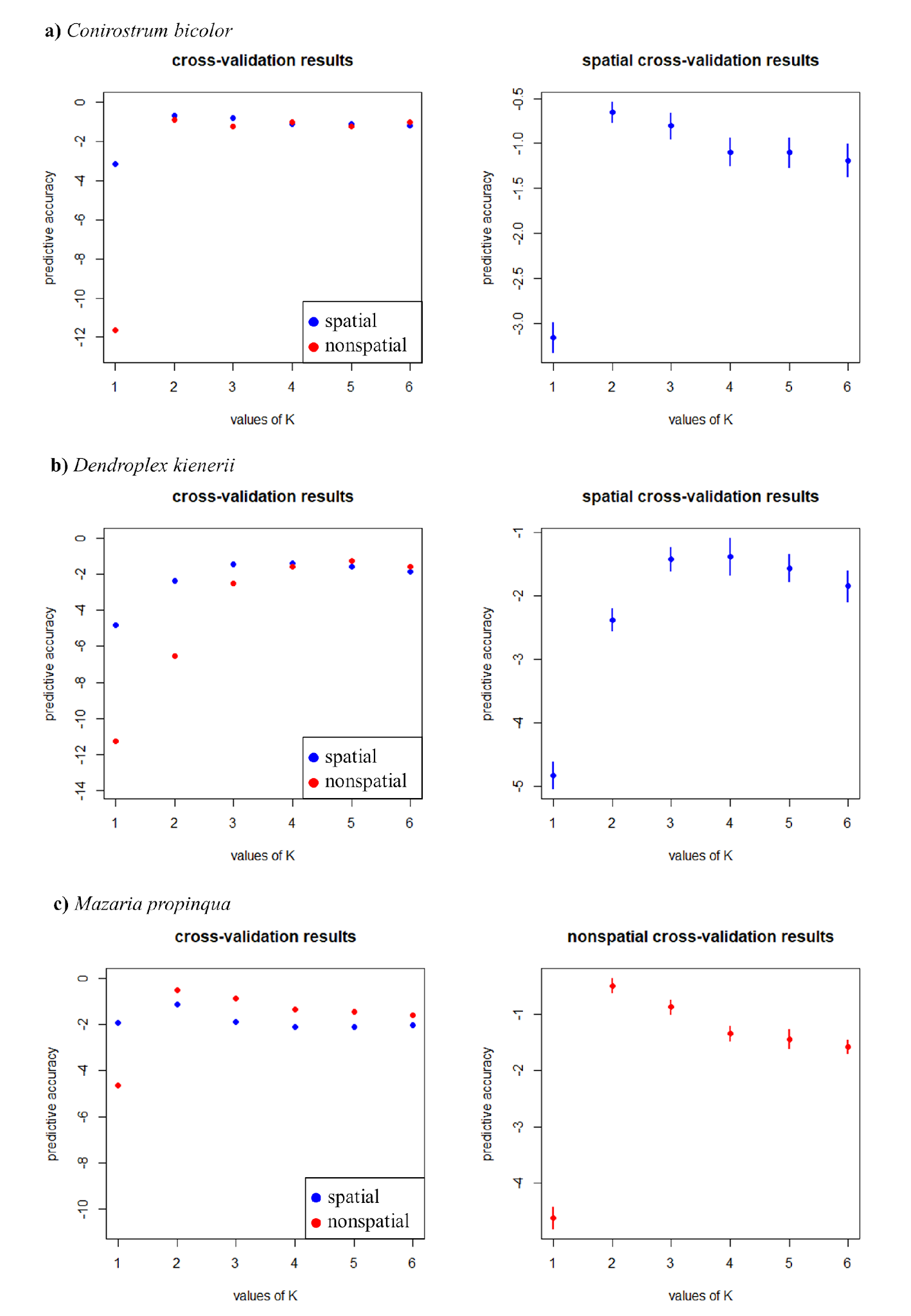

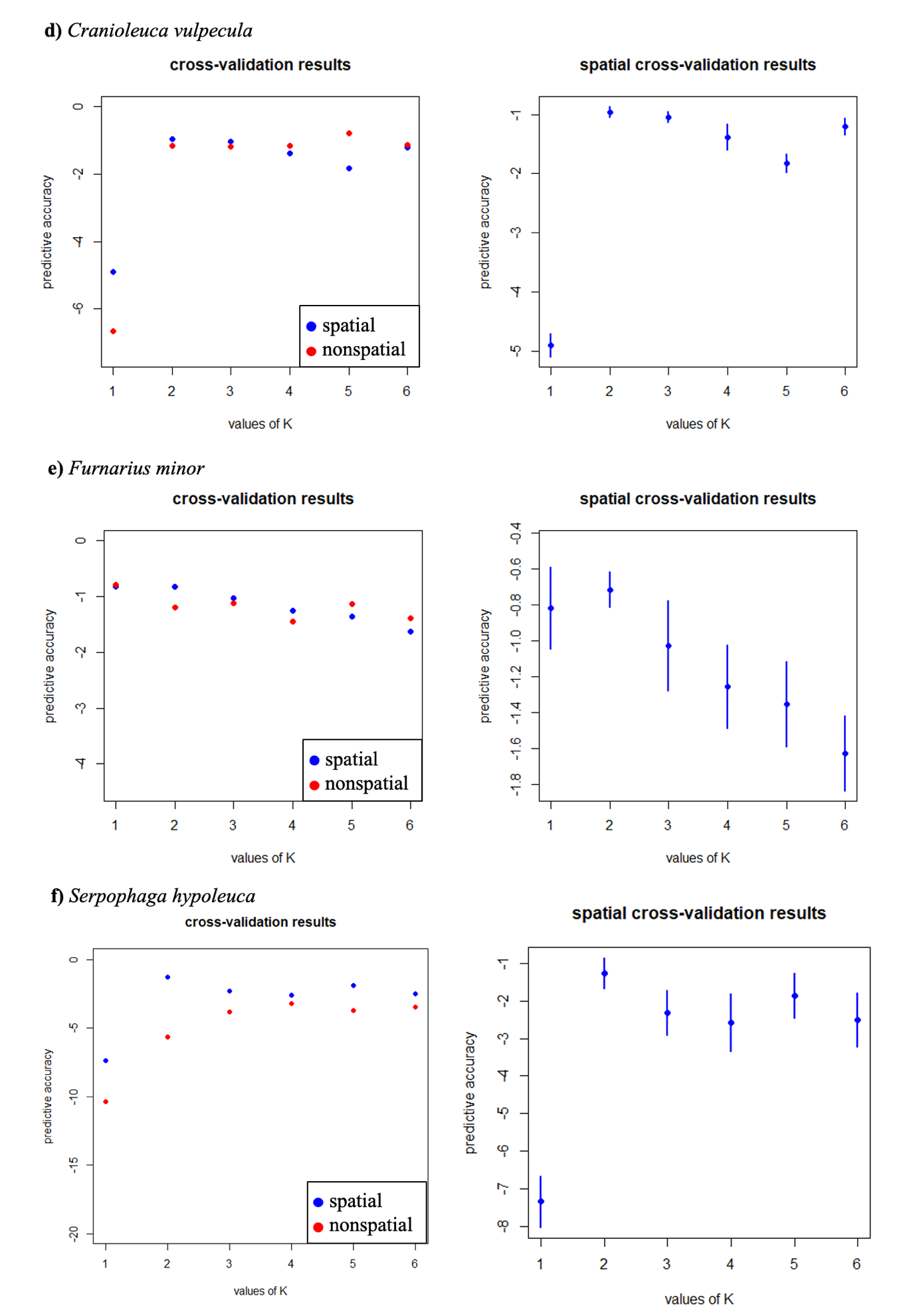

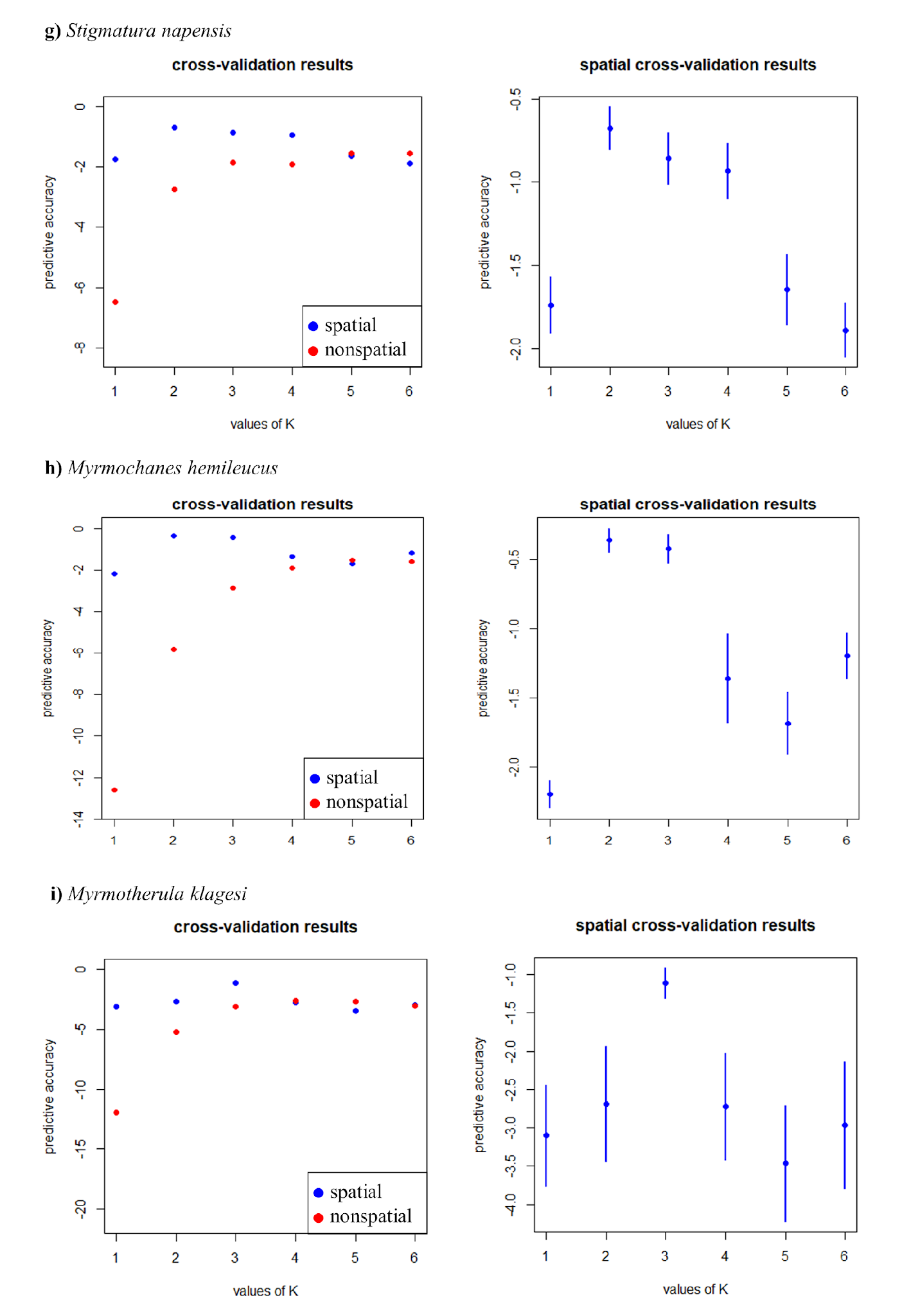
**`**
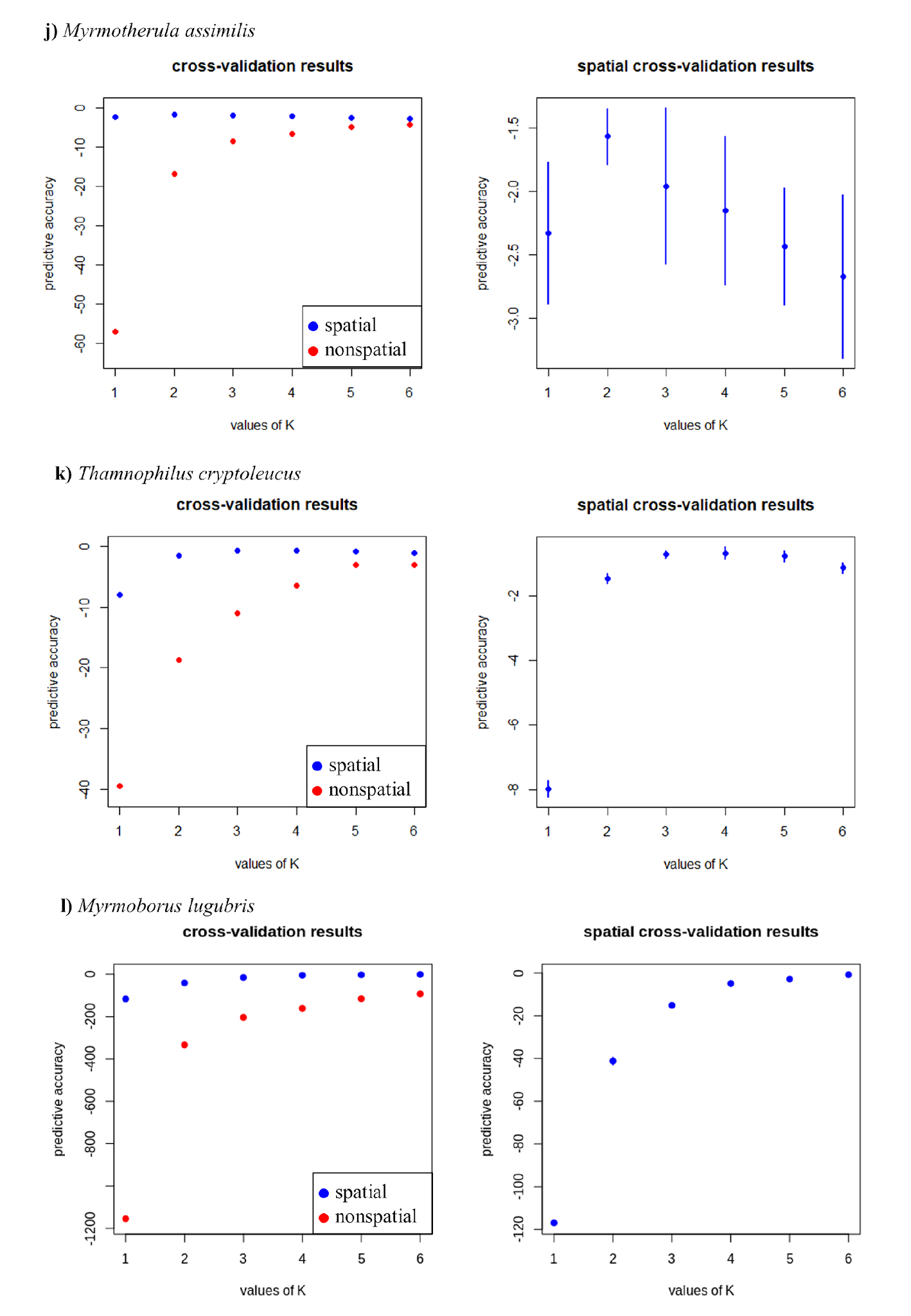

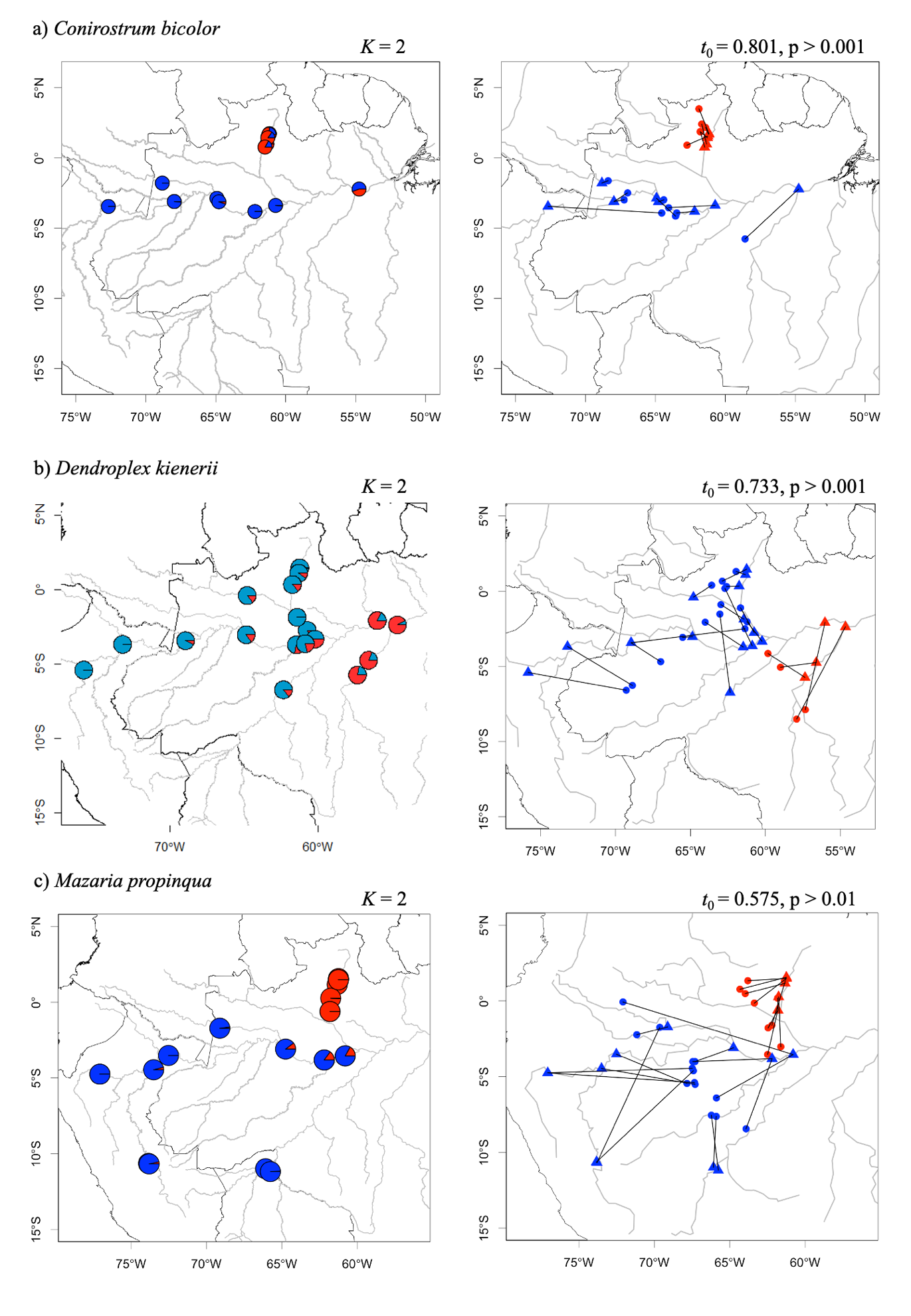

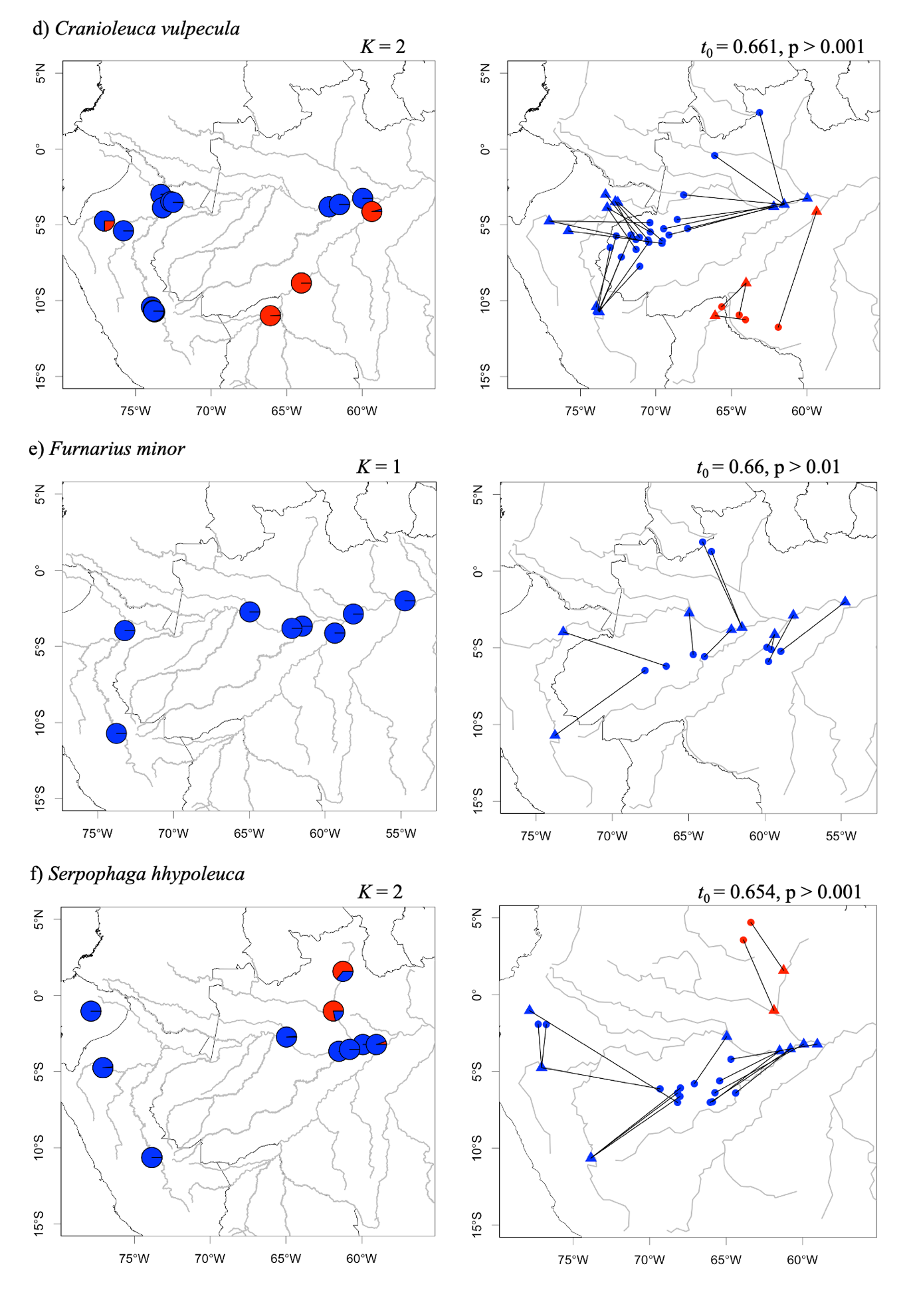

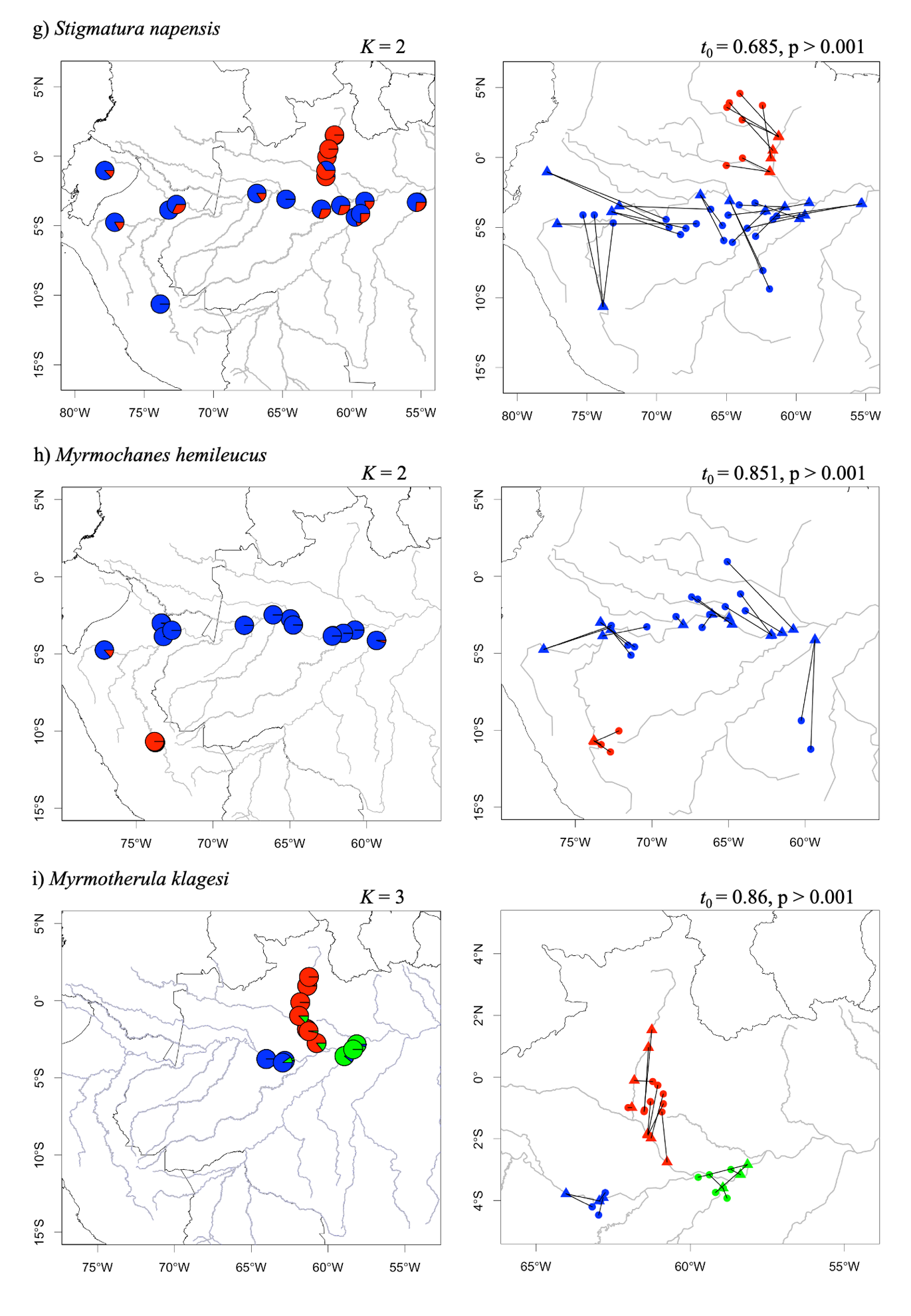

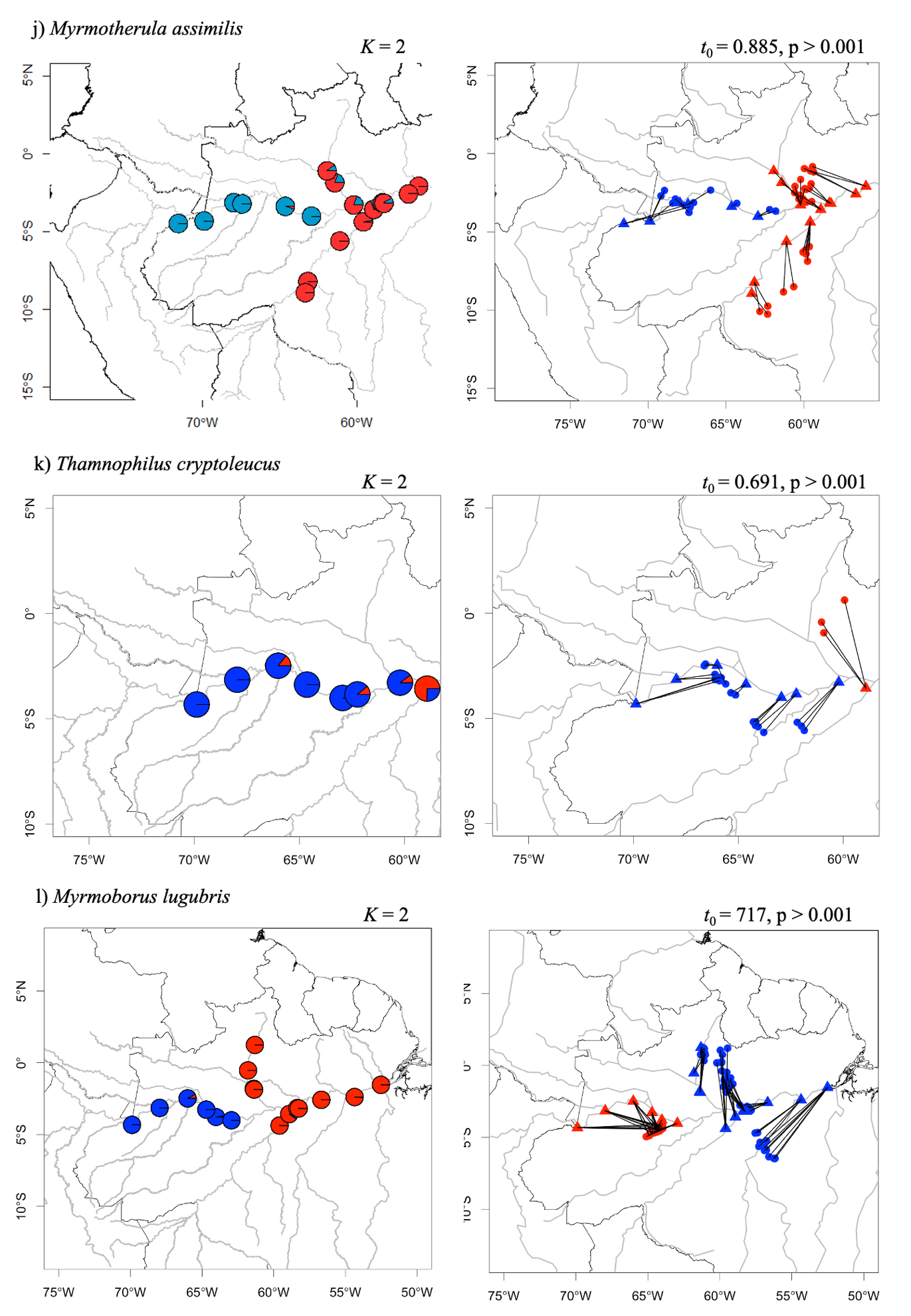
